# Discovery of a Heparan Sulfate Binding Domain in Monkeypox Virus H3 as an Anti-poxviral Drug Target Combining AI and MD Simulations

**DOI:** 10.1101/2024.06.26.600898

**Authors:** Bin Zheng, Meimei Duan, Yifen Huang, Shangchen Wang, Jun Qiu, Zhuojian Lu, Lichao Liu, Guojin Tang, Lin Cheng, Peng Zheng

## Abstract

Viral adhesion to host cells is a critical step in infection for many viruses, including monkeypox virus (MPXV). In MPXV, the H3 protein mediates viral adhesion through its interaction with heparan sulfate (HS), yet the structural details of this interaction have remained elusive. Using AI-based structural prediction tools and molecular dynamics (MD) simulations, we identified a novel, positively charged α-helical domain in H3 that is essential for HS binding. This conserved domain, found across *orthopoxviruses*, was experimentally validated and shown to be critical for viral adhesion, making it an ideal target for antiviral drug development. Targeting this domain, we designed a protein inhibitor, which disrupted the H3-HS interaction, inhibited viral infection *in vitro* and viral replication *in vivo*, offering a promising antiviral candidate. Our findings reveal a novel therapeutic target of MPXV, demonstrating the potential of combination of AI-driven methods and MD simulations to accelerate antiviral drug discovery.

## Introduction

The mpox virus (MPXV), a zoonotic pathogen within the *Orthopoxvirus* genus, has emerged as a global health concern. This genus also includes the Variola virus (VARV), the causative agent of smallpox, and Vaccinia virus (VACV), which has been used as a live vaccine against smallpox^1, 2^ (Fig. 1A). Historically endemic to Central and West Africa, MPXV gained international attention during the global outbreaks in 2022 (clade IIb) and 2024 (clade Ib), leading the World Health Organization (WHO) to declare it a public health emergency of international concern twice within three years^3-5^. Although MPXV has a lower fatality rate than smallpox (2-10% versus 30%), the absence of specific antiviral treatments continues to pose significant health risks^6-11^.

**Fig. 1.**
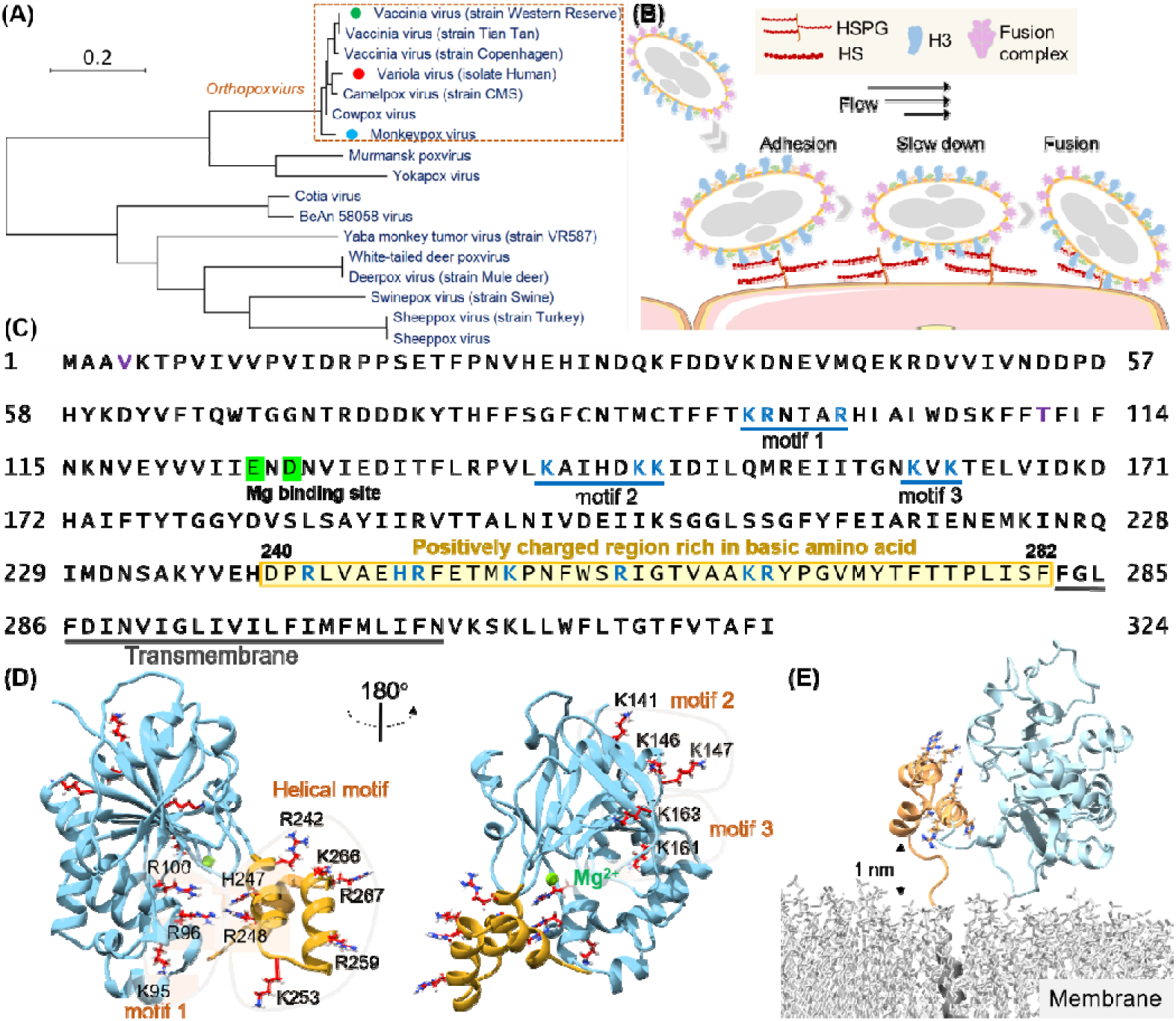
Structural Analysis of the H3 Protein in Monkeypox Virus. (A) Phylogenetic tree depicts the evolutionary relationships of H3 within the *Poxviridae* family, highlighting MPXV (blue circle), VARV (red circle), and VACV (green circle). (B) Schematic of MPXV adhesion to the cell surface. Viral particles bind to cell surface via specific interaction such as between adhesion protein H3 and HS, followed by membrane fusion mediated by the fusion complex, allowing entry into the host cell. (C) The amino acid sequence of MPXV H3 shows the newly discovered helical structure (240-282, highlight in yellow), the Mg(II) (green) binding site and other potential HS binding motifs (blue underlines). (D) AF2 prediction of MPXV H3 structure. The left and right panels show different orientations of the H3 structure (rotated 180°). The blue region corresponds to H3(1-239), which has a homologous crystal structure (VACV H3, PDB code: 5EJ0). The yellow region represents the AF2-predicted structure of H3(240-282), which remains unresolved in the crystal structure of the homologous protein. All potential GAGs-binding motifs are highlighted with a yellow background. (E) MD simulation snapshot of H3 on a DPPC membrane. H3 is anchored to the membrane through its transmembrane region (residues 283-306, in gray).

Viral adhesion to host cells is a critical initial step in the infection process for many viruses. Proteins that mediate these interactions have been identified as essential therapeutic targets, offering promising opportunities for antiviral therapies. In *orthopoxviruse*s such as monkeypox virus (MPXV), the H3 protein plays a pivotal role in facilitating viral adhesion by binding to cell-surface heparan sulfate (HS)^12-14^, a glycosaminoglycan essential for viral entry. H3-specific neutralizing monoclonal antibodies have shown to protective effects for rabbits and immunodeficient mice against lethal poxvirus infections^15^. And H3 is a key component in recent mpox vaccines^16, 17^ Despite the central role of H3 in MPXV infection and protection, the structural details of its specific interaction with HS remain elusive.

Understanding the molecular mechanisms underlying the H3-HS interaction is critical for advancing antiviral drug development. H3 is highly conserved across *poxviruses* (Fig. 1A), including variola and vaccinia viruses, making it a potential target for broad-spectrum therapies. The recent resurgence of MPXV infections has underscored the urgent need for specific antiviral treatments. Given the conserved nature of H3 across all known MPXV lineages, including Clade Ib, which has been linked to the 2024 outbreak, designing therapies that target H3 could offer a robust solution to current and future outbreaks.

However, efforts to resolve the complete structure of H3-HS complex, have been hampered by the dynamic and flexible nature of HS and the tendency of H3 self-cleavage. Classic structural biology techniques, such as X-ray crystallography, have been unable to capture the full interaction between H3 and HS due to these complexities. Therefore, alternative approaches are required to fully elucidate this critical interaction.

To address these challenges, we employed advanced AI-based protein structural prediction and generation tools, including AlphaFold2(AF2) and RFdiffusion, alongside classic molecular dynamics (MD) simulations. These state-of-the-art computational techniques enabled us to identify key HS-binding sites within H3 and to characterize the binding mechanisms of the previously uncharacterized α-helical domain^18-22^. Corroborated by various experimental methods^23-26^, the use of AI-driven tools proved pivotal in overcoming the limitations of traditional structural methods, allowing us to predict and validate the structure-function relationship of the H3-HS interaction. A powerful approach for unraveling complex protein-glycan interactions is demonstrated, offering new pathways for the development of broad-spectrum antiviral therapies^27, 28^.

## Result

### Identification of a Novel Helical Domain for HS Binding in H3

The H3 protein of MPXV is a 324-amino-acid transmembrane protein conserved across all 12 identified *orthopoxviruses*. Previous studies have shown that its extracellular domain (residues 1-282) binds to cell-surface heparan sulfate (HS)^29, 30^, facilitating viral entry in conjunction with other adhesion proteins and the fusion protein system (Fig. 1B)^31^. Inhibiting the HS binding to H3, therefore, represents a promising strategy for developing broad-spectrum antiviral therapies against *orthopoxviruese*. While most structure of the extracellular region of Vaccinia virus H3 (residues 1-240; PDB code: 5EJ0) has been solved, the dynamic and heterogeneous nature of HS as a biopolymer complicates the identification of precise binding sites.

To identify HS binding sites on the H3 protein, we first performed a comprehensive sequence and structural analysis. As a highly negatively charged glycosaminoglycans, it is believed that HS interacts with positive charge residues on proteins by electrostatic interaction. Previous studies have indicated that the positively charged Mg (II) ion in H3 is critical for HS binding. Therefore, we focused on regions rich in basic amino acids—lysine (K), arginine (R), and histidine (H). Given that only the H3 structure from VACV was available^28^, we used AlphaFold2 (AF2) to model the MPXV H3 structure (Fig. S1A), Our analysis revealed three potential HS binding motifs rich in positively charge residues: Motif 1 (K95, R96, R100), Motif 2 (K141, K146, K147), and Motif 3 (K161, K163)^32^, all located within the homologous structure of VACV H3 (Fig. 1C, D).

Interestingly, we also identified a cluster of seven additional positively charged residues (R242, H247, R248, K253, R259, K266, R267) in the protein C-terminal region. This region had not been previously structurally characterized due to its self-cleavage during sample preparation for X-ray diffraction studies, but was retained in all other biochemical and viral assays^28-30^. AF2 predicted these residues to form a three-helical domain with high confidence (pLDDT score: 82.9 for overall structural; 80.6 for the helical domain, 100 for the maximum score), suggesting it may serve as an HS binding site as a structured domain (Fig. 1D). We also tested other models, such as ESMFold (pLDDT:70.6; 50.7) and RoseTTAFold2 (pLDDT: 70.2; 50.7) (Fig. S1B&C) ^33, 34^. They all predict a helical structure. But their results are different in detail and had lower confidence scores. Thus, we used AF2 for further studies. Another concern is that the helical domain near the virus surface. Thus, we modeled the complete structure of H3 in virus using a model DPPC (dipalmitoylphosphatidylcholine) membrane. It showed a dynamic movement between the helical domain and the membrane, with a maximum distance exceeding 1 nm (Fig. 1E, Fig. S1D&E, Supplementary Video 1). Thus, enough space be present for possible HS binding.

Flexible molecular docking and MD simulations identified the exact HS binding sites in the H3 protein.^35, 36^. We used a 20-repeat HS unit with a specific composition for detailed analysis ([IdoA2S-GlcNS6S-IdoA-GlcNS(3,6S)]_5_) (Fig. S2). A 500 ns MD simulation established the equilibrium conformation of H3, and subsequent docking with AutoDock Vina identified four high-scoring conformations for each motif (Fig. 2A, Fig. S3). The docking score calculated by the software reflects the total energy of interactions between molecules. Lower scores (more negative values) indicate more stable binding between the molecules. To address the dynamic nature of HS, we extended our simulations to 3 times additional 1000 ns MD simulations for each conformation. RMSD (Root Mean Square Deviation) analysis demonstrated notably stable binding for HS to both Motif 1 and the newly identified helical domain (Supplementary Video 2), with RMSD values consistently below 4 nm, indicating minimal structural fluctuations during interaction (Fig. 2B).

**Fig. 2.**
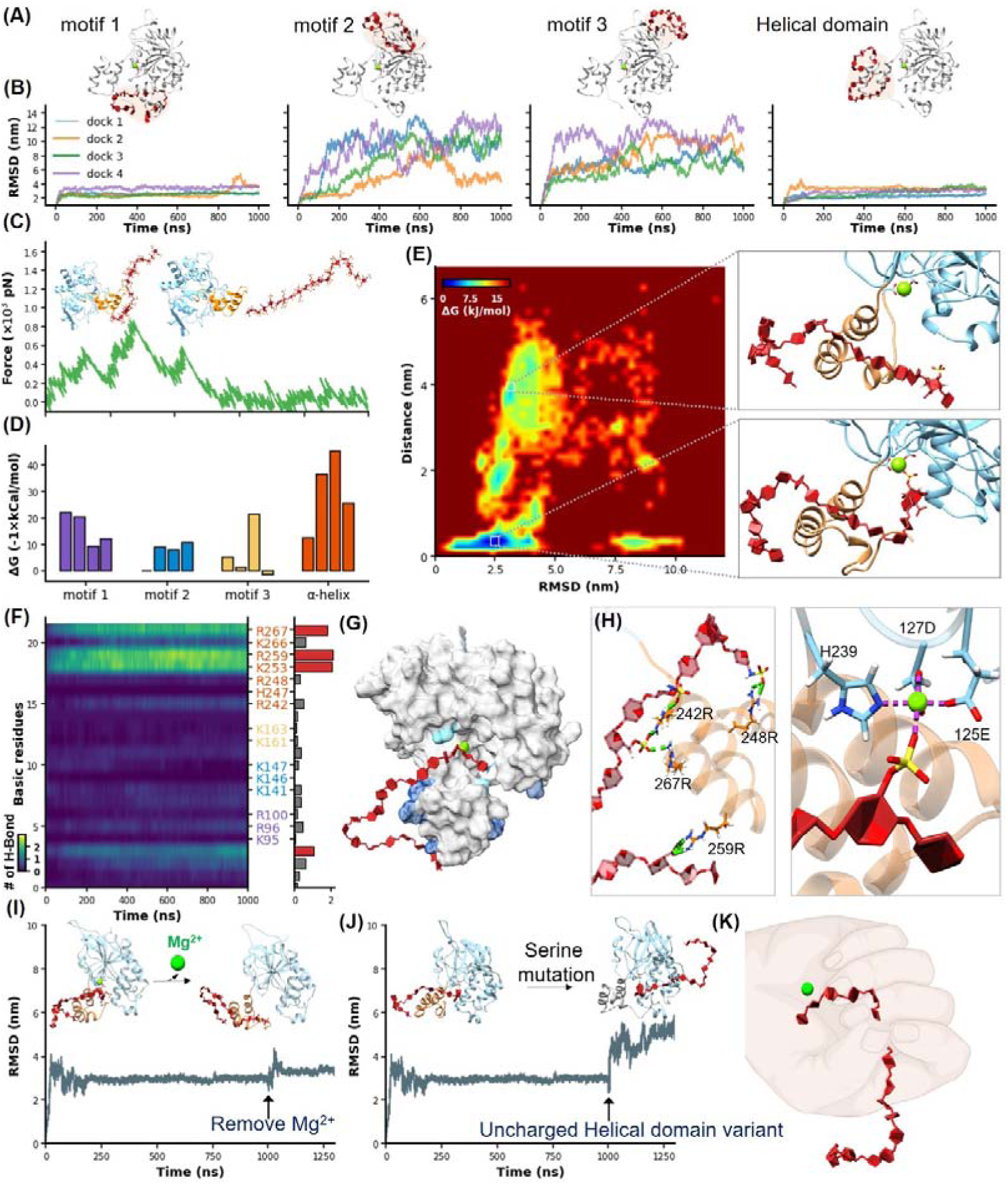
Identification of HS Binding Motifs in H3 by MD Simulation and Docking. (A-B) Cartoons show docking results of HS with Motifs 1, 2, 3, and the helical domain, respectively. Panels (B) show RMSD values from 1µs MD simulations, color-coded to match the configurations in (A). (C) Schematics illustrates the reaction coordinate in umbrella sampling, highlighting HS dissociation from H3 with a green force curve. (D) Histogram shows the binding free energies for HS-H3 interactions in motifs 1, 2, 3, and the helical domain. (E) A free energy landscape map from a 1000 ns REMD simulation shows HS binding configurations to the helical and Mg(□) regions. (F) Panel provides salt bridge formation statistics between HS and H3’s basic amino acids, with a bar chart of average formations. (G) A surface plot shows the frequency of salt bridge formations within H3, with areas of frequent formations in blue. (H) Detailed views of HS-H3 interactions, with the left image showing salt bridges and the right image displaying electrostatic interactions with Mg(□). This panel illustrates the impact on HS binding stability to H3 following the removal of Mg(□) during the simulation. (J) The effects of mutating all basic amino acids in the helical domain on the binding stability of HS are shown. (K) The “palm-binding” model is depicted where HS is secured by the helical “fingers” and interacts with the Mg(□)-bound “palm.”.

Moreover, we quantitatively analyzed the binding free energy between H3 and HS in different conformational states using umbrella sampling (US) techniques. A harmonic potential was applied to HS and gradually pulled away from H3, generating a series of reaction coordinates (Fig. 2C, Fig. S4, Supplementary Video 3). The green force-extension curve represents the force during this stretching process. This curve, along with the corresponding PMF (Potential of Mean Force) variation shown in Figure S4, demonstrated the free energy changes throughout the umbrella sampling process, revealing that the helical domain possesses the highest binding affinity with a ΔG of -45 kCal/mol (Fig. 2D).

Previous studies have suggested the critical role of the Mg(II) ion in stabilizing HS binding, where its removal reduces binding efficiency^29^. Indeed, the α-helical domain is very close to the Mg(□)-bound region in H3. Notably, these two parts appear to form a distinct cavity-like binding pocket for HS (Fig. 1E, Fig. 2E). Thus, we conducted a 1000 ns REMD (Replica Exchange Molecular Dynamics) simulation of the H3-HS complex to further understand the binding dynamics. REMD, an advanced sampling technique, helps overcome the kinetic barriers typically encountered in conformational transitions by utilizing a series of temperature-controlled replicas to enhance the exploration of the conformational landscape^37^. For this study, we set up 64 replicas over a temperature range from 310K to 387K, cumulatively simulating for 64 µs. This comprehensive simulation captured the HS molecule progressively binding deeper into a cavity formed by the α-helical domain, eventually interacting with the Mg (II) ion as hypothesized (Fig. 2E, Supplementary Video 4).

The free energy landscape, mapped out from these simulations, shows HS transitioning from a stable interaction with the helical domain across an energy barrier into the Mg (II)-enhanced cavity. This dual interaction—first with the helical domain and then with the Mg(II) site—was consistently observed to stabilize the complex further, evidenced by a significantly lower binding free energy of -67 kCal/mol (Fig. S4)^38^. These results underscore the dynamic nature of the H3-HS interaction and validate our model of sequential binding, which could be critical for designing inhibitors that target these specific interactions.

Given the crucial role of electrostatic interactions in the HS-H3 binding process, we conducted an exhaustive analysis of salt bridge formations across all MD simulation trajectories. This analysis focused on the formation of salt bridges over time between the basic amino acids (R, K, H) of H3 and HS, with brighter areas indicating more frequent formation of salt bridges and robust interaction (Fig. 2F, Fig. S5). A bar graph accompanied these maps, quantifying the average number of salt bridges each amino acid formed during the simulations, further illustrating the intensity and frequency of these interactions. Our findings demonstrated a high affinity of HS for the helical domain of H3, with basic amino acids in this region forming more salt bridges compared to other parts of the protein. The surface hotspot map of H3 (Fig. 2G) visually highlighted these interactions, with varying shades of blue representing the frequency of salt bridge formation, emphasizing their distribution and intensity. In the helical domain, a focus view showed that the 1st, 7th, 9th, and 18th sugar units of HS formed salt bridges with R248, R242, R267, and R259 of H3, respectively (Fig. 2H, left), while the sulfate group on the 2nd sugar side chain of HS bound to Mg(II) (Fig. 2H, right).

To further prove these findings, we performed two control experiments. First, we removed the Mg(□) ion during MD simulation. According to the results of umbrella sampling, we extended the simulation of the most stable H3-HS binding conformation to 1000 ns, then removed the Mg(□) ion and continued the simulations. The resulting increase in RMSD fluctuations confirmed the stabilizing role of Mg(□) in the binding process (Fig. 2I, Supplementary Video 5). Second, we mutated all seven positively charged amino acids in the helical domain to serine, and name it H3 (uncharged). While AF2 predicted that the helical structure would remain intact (Fig. S6), a significant alteration in HS binding dynamics was observed, supporting the critical role of these charged residues in stabilizing the interaction (Fig. 2J, Supplementary Video 6).

Our interaction model revealed that the α-helical domain of H3 functions like a thumb, guiding HS into a stable binding position, while the Mg(□) region acts like a palm, securing HS in place. This dynamic interaction allows HS to transition from an initial surface-level binding to being deeply anchored within the protein cavity, illustrating a complex yet well-organized binding process (Fig. 2K).

Bioinformatic analysis of the H3 protein across the *Poxviridae* family highlights its evolutionary conservation and its significance in HS binding. Given the crucial role of electrostatic interactions in the H3-HS binding process, we analyzed the charge distribution within the helical domain. To date, H3 proteins have been discovered in 66 of 118 known *Poxviridae* species. Using NCBI’s BLAST, we retrieved sequences of these 66 H3 proteins and performed multiple sequence aligment^39^. A heatmap detailing the side-chain charge distribution at a physiological pH of 6.5—mimicking the microenvironment of H3-HS interaction—revealed a significant concentration of positive charges at the helical domain (Fig. 3A). Furthermore, structural predictions using AF2 showed that all these 66 sequences generally share a similar architecture to MPXV H3, particularly in the presence of the α-helical domain (Fig. S7).

**Fig. 3.**
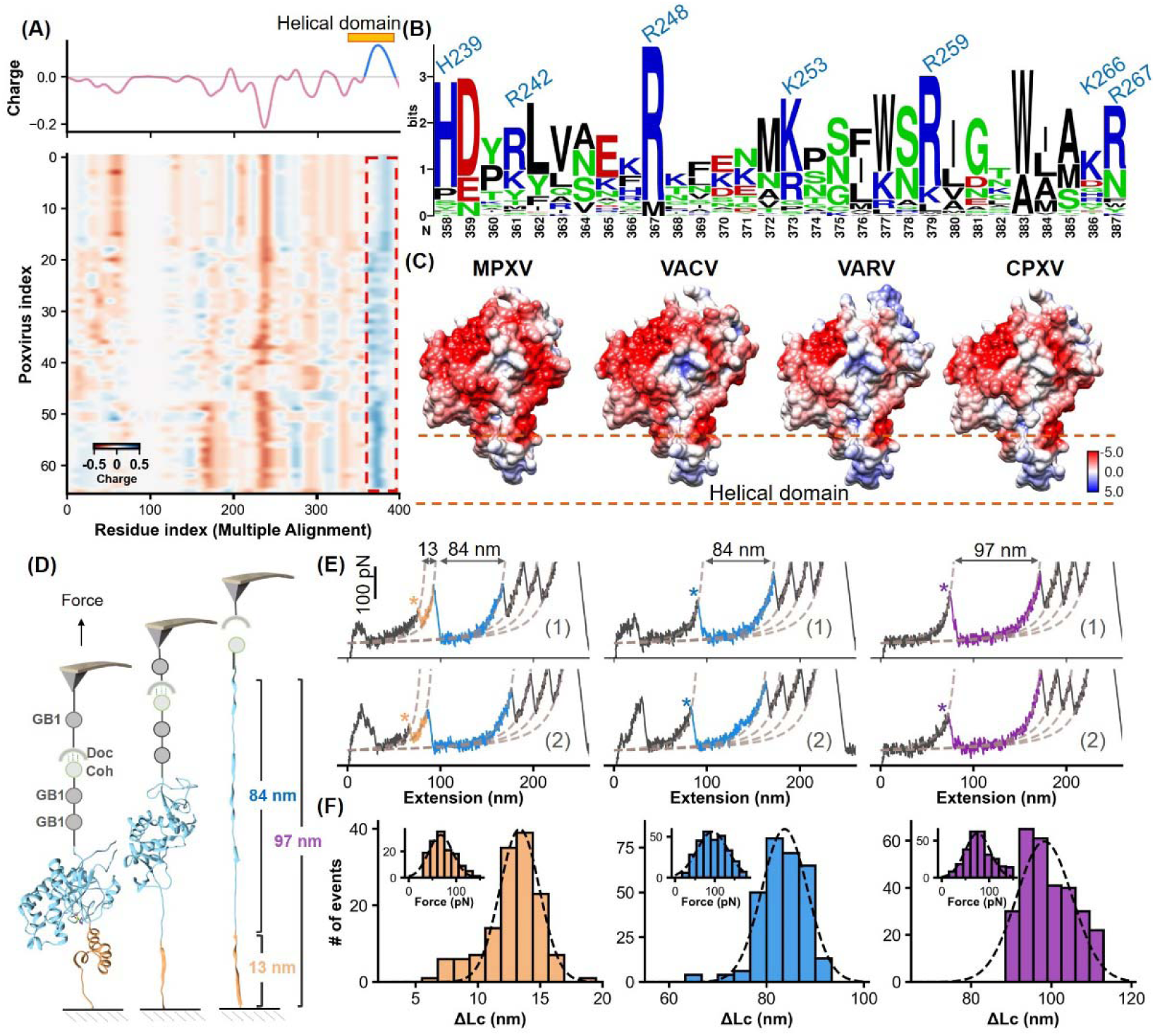
Charge Characteristics and Structural Analysis of H3 Protein. (A) The heatmap shows the amino acid charge distribution in the H3 protein across 66 *Poxviridae* viruses, following multiple sequence alignment. Blue indicates areas with more positive charges, and red indicates more negative charges. The accompanying curve shows the average charge of all amino acids in *Poxviridae* H3. (B) Logo plot of the amino acid sequence of the helical region, highlighting the conservation of basic amino acids at specific positions. (C) The surface charge analysis of H3 from the MPXV, VACV, VARV, and cowpox viruses (CPXV) with the helical domain showing a significantly positive charge. (D) Schematic of the single-molecule force spectroscopy unfolding experiment on H3, illustrating the unfolding process of the helical domain (yellow) followed by the main body (blue). (E) Representative curves of H3 unfolding, color-coded to show the helical region (yellow), main body (blue), and full-length (purple) unfolding. (F) Histograms depict the force spectroscopy signals for helical domain, main body, and full-length unfolding, with ΔLc statistics provided. The inset shows a Gaussian fit of unfolding forces.

Notably, this domain consistently exhibited an accumulation of basic amino acids such as lysine, arginine, and histidine, which are essential for binding the negatively charged HS. These amino acids, including residues identified as 358, 361, 367, 373, 379, 386, and 387 in *Poxviridae* H3 (global position in alignment), corresponding to residues H239, R242, R248, K253, R259, K266, R267 in MPXV H3, show high conservation (Fig. 3B, Fig. S8). Surface charge analysis of H3 proteins from four *orthopoxviruses*, including MPXV, VACV, VARV, and CPXV confirmed the predominance of strong positive charges in their α-helical domains (Fig. 3C), crucial for effective binding to HS. This consistent feature across different viruses underscores the evolutionary importance of the helical domain, suggesting a universal mechanism in orthopoxviral adhesion that could be targeted in antiviral strategy.

### Experimental Confirmation of the Structure and Function of α-Helical Domain

To validate the computational predictions, we first conducted experiments using atomic force microscopy-based single-molecule force spectroscopy (AFM-SMFS). This technique, widely used for studying protein (un)folding, allowed us to directly assess the structure and stability of the α-helical domain within the H3 protein^40-42^. We engineered the H3 construct with two GB1 domains, each 18 nm in length upon unfolding, and one Cohesin module for precise measurement (Fig. 3D)^43-45^. These proteins are immobilized on the surface by click chemistry and enzymatic ligations (Fig. S9)^46-48^. During the experiment, the coated AFM tip with GB1 and Dockerin initiated a Coh-Doc interaction, producing a characteristic sawtooth-like force-extension curve as it was stretched (Fig. 3D&E). Notably, two peaks corresponding to the unfolding of the α-helical domain were observed. The first peak showed a contour length increment (ΔLc) of 13 nm, closely matching the theoretical unfolding length for the 42 amino acid-long α-helical domain of H3 (residue 240-282, 42aa*0.36 nm/aa-2.6 nm)^49^. The measured unfolding force was 67.4 ± 2.1 pN (mean ± SEM, *n*=139), while the second peak indicated the unfolding of the remaining structure of H3, with a ΔLc of 84 nm (Fig. 3F). The total unfolding, represented by a cumulative ΔLc of 97 nm, confirmed the structured and stable nature of the α-helical domain within H3.

We further explored the functional role of the α-helical domain in mediating H3’s binding to HS using AFM on live cells^50, 51^. The H3 protein was linked to the AFM tip and approached cultured CHO-K1 cells, which express HS, mounted on a Petri dish. Inverted microscopy was used to precisely control the positioning of the AFM tip. Upon contact, the interaction between H3 and HS was initiated, and the binding event was monitored by retracting the tip to generate a force-extension curve. A distinct peak, corresponding to H3-HS interaction, was observed with a dissociation force of 33.7 ± 0.2 pN and a binding probability of 43% (Fig. 4A, B). Repeated trials across different cell surface areas produced a detailed force map, quantifying the distribution of HS unbinding forces and further confirming the critical role of the helical domain in viral adhesion (Fig. 4C, D).

**Fig. 4.**
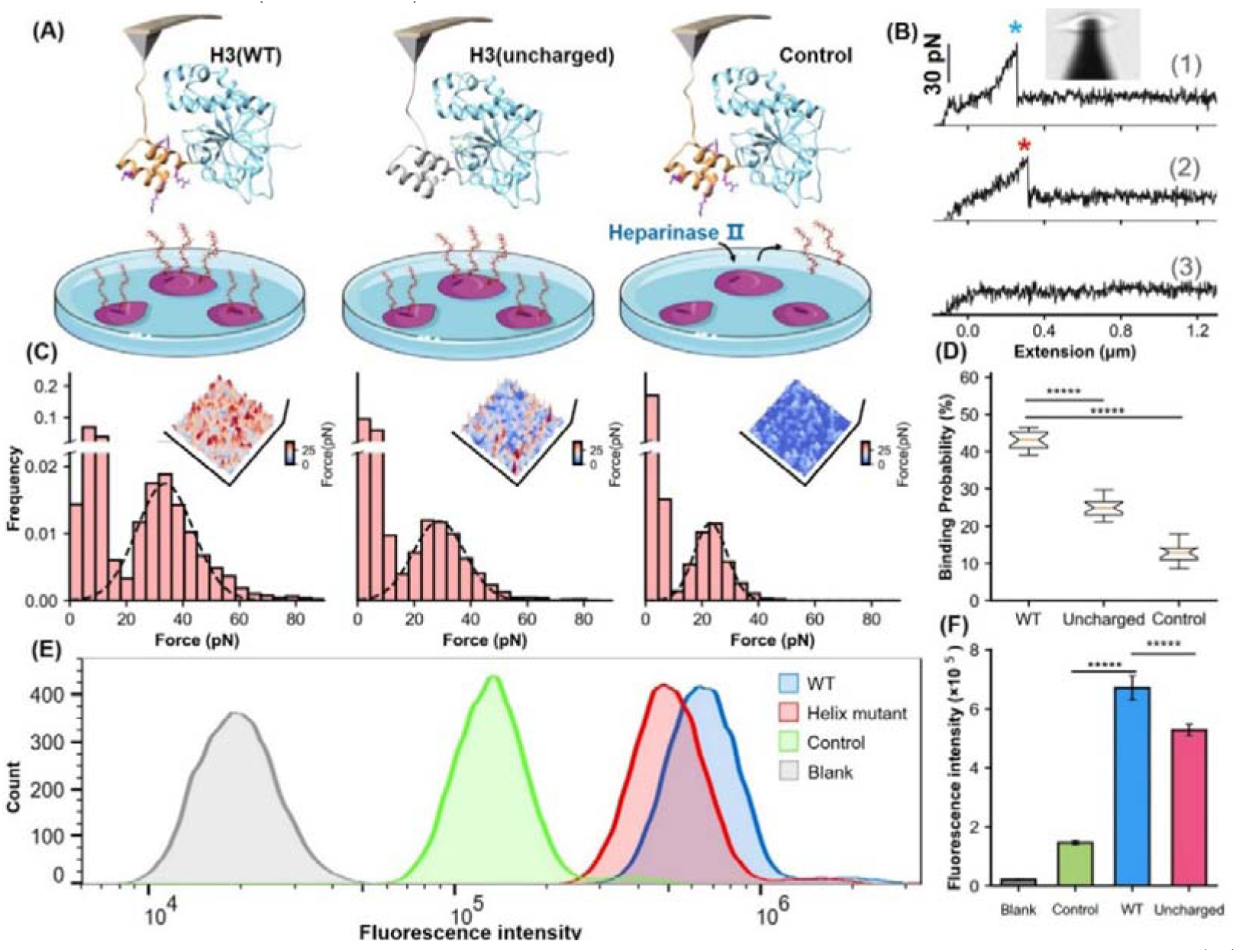
Analysis of the Helical Domain’s Interaction with HS at Cellular Level. (A) Schematic of the cell force spectroscopy experiment setup shows three scenarios: wild-type H3 on an AFM tip interacting with HS on CHO-K1 cells, mutation of all basic amino acids in the H3 helical region to serine, and cells treated with HS hydrolase to remove surface HS before testing. (B) Force-extension curves depict interactions for the wild type, mutant, and control groups, marked with blue and red asterisks for dissociation events. An inset shows the optical microscope positioning the AFM probe. (C) Histograms of dissociation signals comparing the wild type, mutant, and control groups, with an inset detailing the surface distribution of dissociation forces. (D) Statistical graph showing binding probabilities for different groups, highlighting significant differences determined by t-tests (p<1e-5). (E) FCM results illustrate interactions of wild-type (WT) and Uncharged H3 fused with eGFP with CHO-K1 cells, alongside GFP control (green) and cell-only control (Blank, grey). (F) Statistical analysis of FCM data, showing significant differences between groups as determined by t-tests. *****, P<1e-5

Moreover, we performed mutagenesis studies on the α-helical domain by replacing all positively charged residues with serine. AFM experiments on this variant H3(uncharged) showed a detectable force peak with a lower unbinding force of 28.8 ± 0.2 pN, compared to 33.7 ± 0.3 pN for the wild type—representing a 14% reduction (ΔForce > 2*SEM, 95% CI)—and a significant decreased HS binding probability from 43% to 25% (Fig. 4C, D). These results suggest that the basic amino acids in the helical domain are crucial for HS binding. To confirm the specificity of the interaction, CHO-K1 cells were treated with heparinase II, an enzyme that degrades HS. This treatment further reduced the dissociation force to 23.0 pN and the binding probability to 12.6%, confirming that the observed interactions were specific to the HS-H3 complex.

Further validation was carried out using flow cytometry (FCM). We fused eGFP, a fluorescent reporter protein, to the C-terminus of both the H3(WT) and the H3(uncharged). Following incubation of these fusion proteins with cells, FCM analysis showed a significant reduction in mean fluorescence intensity in cells treated with H3(uncharged) compared to H3(WT) (Fig. 4E). Statistical analysis of multiple experiments highlighted a significant difference between the two (Fig. 4F). These findings, along with the results from MD simulations and AFM force spectroscopy, underscore the importance of the positively charged amino acids in the α-helical domain of H3 for efficient HS binding.

### The Helical Domain Serves as a Target for H3 Infection Inhibition

Targeting the α-helical domain of H3 as H3 binding sites, we *de novo* designed a series of inhibitors. Using deep-learning model RFdiffusion, we designed a series of inhibitors for the H3-HS interaction and named it AI-PoxBlock (Fig. 5A). Sequence recovery for each series was performed on 1000 backbones using ProteinMPNN, generating ten sequences per backbone. These designs were further validated through complex structure predictions using AF2-multimer, and candidates were selected based on PAE (Predicted Aligned Error) values below 7 and iPTM (interface predicted Template Modelling) scores above 0.9. The predicted structures of these candidates also exhibited RMSD values of less than 2 nm when compared to the RFdiffusion-generated structures (Fig. S10). To assess the binding capabilities of these inhibitors, we conducted 500 ns MD simulations, analyzing RMSD trajectories. Five inhibitors—AI-PoxBlock302, 602, 614, 723, and 761—demonstrated stable RMSD values under 0.4 nm, confirming their potential for stable binding to the helical domain of H3 (Fig. S11, S12, Supplementary Video 7).

**Fig. 5.**
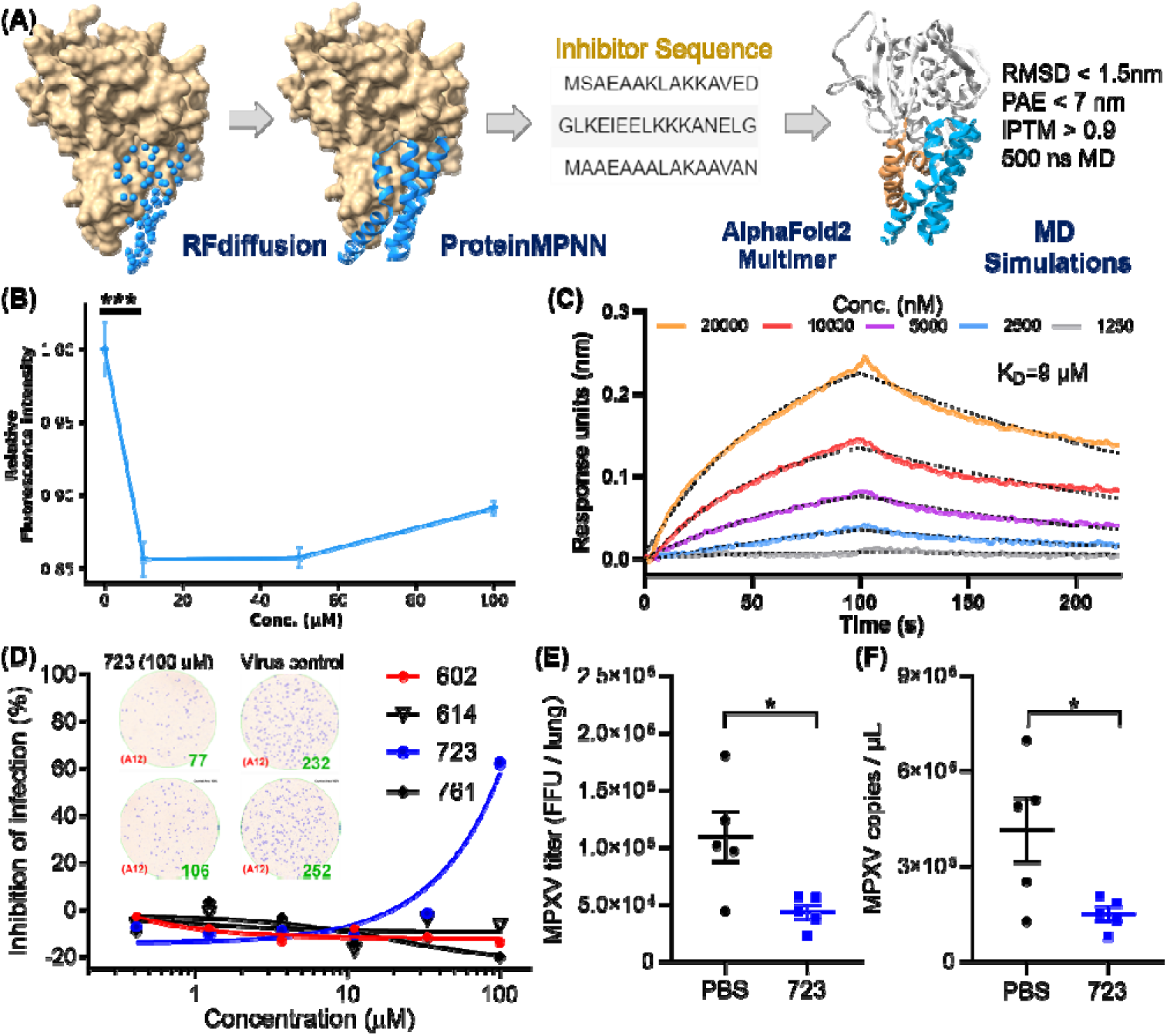
Inhibitor Design Targeting the H3-HS Interaction and Efficacy of AI-PoxBlock723. (A) Diagrams depict protein inhibitors designed to target the H3 helical region, created using RFdiffusion. Sequences capable of folding into the target scaffold structures were generated using ProteinMPNN, and were validated through AF2, followed by 500 ns MD simulations for structural stability and interaction scoring. (B) FCM analysis demonstrates the inhibitory effect of AI-Poxblock723 at various concentrations (0, 10, 50 and 100 μM, *x* axis). The control group consisted of 2 μM H3-eGFP without the inhibitor, with the relative fluorescence intensity normalized to 1. (C) BLI confirms direct interaction between AI-Poxblock723 and the H3 helical domain. (D) Graphs display the inhibitory effect of the indicated AI-Poxblocks on MPXV infection of Vero E6 cells, with quantitative analysis of virus-infected foci provided on the right. (E-F) BALB/c mice were infected intranasally with 4 × 10^5^ FFU MPXV and treated with single dose of AI-PoxBlock723 (10 mg/kg) or PBS immediately after challenge (*n* = 5). Infectious MPXV particles and MPXV viral loads in mice lungs at 4 dpi were determined by focus-forming assay (E) and qPCR (F).

Although AI-PoxBlock302 could not be successfully expressed, the remaining four inhibitors underwent further testing in FCM to evaluate their efficacy in inhibiting H3 binding to CHO-K1 cell surfaces. AI-PoxBlock723 stood out, significantly reducing H3 binding at a concentration of 10 μM, while the other inhibitors displayed no tendency to inhibit H3 binding to HS on cell surfaces (Fig. 5B, Fig. S13). Further analysis using biolayer interferometry (BLI) confirmed that AI-PoxBlock723 binds to H3 with an equilibrium dissociation constant (*K*_D_) of 9 μM (Fig. 5C). To ensure specificity, we also assessed the interaction between AI-PoxBlock723 and a truncated version of H3 (residues 1-239), lacking the α-helical domain. BLI results indicated a marked reduction in binding, reinforcing the specificity of AI-PoxBlock723 for the helical domain (Fig. S14). Circular Dichroism spectroscopy also confirmed that AI-PoxBlock723 predominantly consists of α-helices, consistent with its design specifications (Fig. S15).

To evaluate the antiviral activity of AI-PoxBlocks, the inhibitors were serially diluted and incubated with the clade II MPXV isolate SZTH42 before infecting Vero E6 cells. After overnight culture, virus infection foci were determined as previously described.^52^ Notably, AI-PoxBlock723 showed an antiviral effect with the half maximal inhibitory concentration (IC_50_) of 88.6 μM, whereas other AI-PoxBlocks exhibited no inhibitory efficiency (Fig. 5D). Given the high conservation of the H3 helical domain across the four *orthopoxviruses* pathogenic to humans (Fig. S16), we also assessed AI-PoxBlock activity against VACV infection. Consistently, AI-PoxBlock723 is also effective for inhibiting VACA infection in vitro with a similar IC50 of 86.7 μM (Fig. S17), suggesting the helical domain as a universal antiviral target for *orthopoxvirus*.

To evaluate the efficacy of AI-PoxBlock723 in vivo, groups of BALB/c mice were challenged with 4 × 10^5^ FFU (focus forming unit) MPXV/SZTH42 as we previously reported^53^. Mice were administered intraperitoneally with single dose of AI-PoxBlock723 (10 mg/kg) or PBS immediately after MPXV challenge, and were sacrificed 4 days post-infection (dpi) for the determination of pulmonary MPXV titers. The MPXV infectious particles and genome copies in lungs of the AI-PoxBlock723 treated mice were significantly lower than that in the control mice (Fig. 5E-F, Fig. S18).

These results demonstrate the efficacy of the designed inhibitors targeting the H3 helical domain and validate the crucial role of this domain in viral adhesion, providing a promising foundation for the development of novel antiviral agents.

## Discussion

Our study provides new mechanistic insights into MPXV infection by identifying the α-helical domain of the H3 protein as a critical mediator of heparan sulfate (HS) binding. This domain, previously uncharacterized, plays a pivotal role in viral adhesion to host cells, a key step in MPXV pathogenesis. Using a combination of AI-based tools and molecular dynamics (MD) simulations, we successfully identified the specific interaction sites between H3 and HS, offering a novel target for antiviral drug design^54^.

Previous studies have highlighted the importance of viral adhesion proteins in *orthopoxvirus* infection, but the specific mechanisms underlying the H3-HS interaction remained unresolved. Our findings provide the first detailed structural insight into this interaction, confirming the importance of the electrostatic nature of H3-HS binding. These results align with prior reports indicating the role of HS in viral entry, but advance the field by identifying specific residues within the α-helical domain that are crucial for binding.

One of the strengths of this study lies in the application of advanced AI-based tools, such as AlphaFold2 and RFdiffusion, alongside MD simulations. These cutting-edge computational approaches allowed us to overcome the limitations imposed by the dynamic and flexible nature of HS. In addition, HS serves as a cellular ligand for various viruses^55-57^, underscoring the importance of studying its interactions with viral adhesion proteins. By predicting the structure-function relationship of the H3-HS interaction, we were able to pinpoint key binding residues and mechanisms that were previously inaccessible. This demonstrates the growing potential of AI in structural biology, particularly for the complex protein-glycan interactions^56, 58,59^.

The identification of the α-helical domain in H3 as a key player in HS binding offers a promising new target for antiviral drug development^17, 60-67^. Given the conservation of this domain across *orthopoxviruses*, including variola and vaccinia viruses, targeting this region could lead to the development of broad-spectrum antivirals effective against multiple *orthopoxviruses*^68^. Our study highlights the power of AI-driven drug design, as demonstrated by the promising results of AI-PoxBlock723, which showed efficacy in inhibiting viral infection. Further optimization and testing of these inhibitors in clinical models will be critical steps toward developing effective antiviral therapies. For example, conjugate the peptides with carrier molecules, such as liposomes, nanoparticles, or dendrimers, which can protect the peptides from immune detection and improve their delivery to target cells.

Looking ahead, the methodologies and findings from this study are likely to catalyze further investigations into viral adhesion mechanisms, not only in *orthopoxviruses* but also across other viral families that utilize similar adhesion strategies. Although challenges remain, we hope our discovery will attract structural biologists to resolve this domain’s structure in H3, and even the structure of H3-HS complex, to further enhance drug design efficiency for this target. Although this domain itself cannot express, we chemically synthesized it (residues 240-279). CD measurement on this fragment indeed showed an α-helical structure (Fig. S19). As AI-driven approaches continue to evolve, they are poised to play a transformative role in advancing our understanding of viral pathogenesis and revolutionizing the future of drug discovery.

## Acknowledgments

We appreciate the inspiring comments from previous anonymous reviewers. We acknowledge the funding support from the National Natural Science Foundation of China (22222703, 22477058), the Fundamental Research Funds for the Central Universities (020514380335), the Natural Science Foundation of Jiangsu Province (BK20202004), Shenzhen Medical Research Funds (B2302052). The numerical calculations in this work have been done on the computing facilities in the High-Performance Computing Center (HPCC) of Nanjing University.

## Author contributions

Conceptualization: PZ

Investigation: BZ, MD, YX, JQ, SW, RG, GT, LC

Funding acquisition: PZ, LC

Project administration & Supervision: PZ

Writing – original draft: PZ, BZ, LC

## Data and materials availability

All data are available in the main text or the supplementary materials

## Supplementary Materials

includes supplementary materials and methods, notes, figures and videos.

