## Supplementary material for "Discovery of a Heparan Sulfate Binding Domain in Monkeypox Virus H3 as an Anti-poxviral Drug Target Combining AI and MD Simulations": supplmentary information

**SUPPLEMENTARY MATERIALS**

**Materials and Methods**

**Protein expression and purification**

All genes were ordered from General Biosystems. H3(1-282) was constructed into the pCold-TF-tev vector using *BamH*Ⅰ-*Bgl*Ⅱ-*Kpn*Ⅰ three-restriction enzyme system and a Strep-Tag II was added to the C-terminus for protein purification. eGFP was ligated to the C-terminus of H3(1-282) by restriction enzyme system. Mutations in the basic amino acids of the helical domain were introduced by site-directed PCR mutagenesis. The pCold-TF-tev-POI-strep tag construct was transformed into BL21(DE3), and single colonies were picked and cultured overnight at 37°C in LB medium with ampicillin. The expanded culture was then transferred to 800 mL of ampicillin LB and continued to be incubated at 37°C with shaking for 2 hours. When the OD_600_ of the culture reached 0.6-0.8, it was cooled to 15°C in an ice-water bath. Then, IPTG was added to a final concentration of 0.5 mM and incubated overnight at 15°C on a shaker.

The next day, the cells were collected by centrifugation at 8000 rpm, the supernatant was discarded, and the cell pellet was resuspended in buffer (50 mM Tris-HCl, 150 mM NaCl, pH 7.4). The cell suspension was then lysed with a homogenizer for 3 minutes, and the lysate was centrifuged at 18000 rpm to separate the precipitate. TEV protease was added to the supernatant, along with EDTA and DTT to a final concentration of 1mM each, and incubated at 30°C for 3 hours to remove the TF tag. The digested product was subjected to another high-speed centrifugation at 18000 rpm, and the supernatant was collected. The supernatant was then purified using STarm Sreptactin. The purity of the obtained protein was verified by SDS-PAGE, and the final purity of the H3 protein exceeded 60%.

The proteins used for AFM force spectroscopy were sequentially constructed into the pQE80L vector using the same restriction enzyme system, resulting in pQE80L-coh-(GB1)2-H3(1-282)-strep tag Ⅱ-NAL. The protein expression method was similar to the one mentioned above. The difference was that after adding IPTG to a final concentration of 0.5 mM, the culture was incubated overnight at 18°C. After cell lysis and centrifugal removal of the precipitate, the proteins were purified using STarm Sreptactin. The eluate was exchanged into AFM working buffer (50 mM Tris-HCl, 150 mM NaCl, pH 7.4) using an ultrafiltration tube and then used for AFM force spectroscopy experiments.

*Oa*AEP1(C247A) is cysteine 247 to alanine mutant of asparaginyl endoproteases 1 from *oldenlandia affinis*, abbreviated as AEP here. ELP is the elastin-like polypeptides (2). Their expression and purification protocols can be found in references. The TEV protease was expressed in the pCold-His6-ProS2 vector (the original tev protease cleavage site in the vector was removed by PCR). The expression was induced with a final concentration of 0.1mM IPTG and incubated overnight at 15°C in a constant temperature shaker. The His6-ProS2-TEV was purified using Ni-NTA affinity chromatography.

All H3 inhibitors designed via RFdiffusion were constructed into the pET30a vector, with C-terminal fusion of strep tag and His8 tag for protein purification and subsequent BLI immobilization. After transforming pET30-Inhibitor-strep tag Ⅱ-His8 into BL21(DE3), the culture was grown at 37°C in kanamycin-resistant LB medium for 2 hours until the OD_600_ reached 0.6-0.8, then IPTG was added to a final concentration of 0.5 mM and the culture was incubated overnight at 18°C. The subsequent purification steps were as previously described, using STarm Sreptactin. The purified protein was exchanged into PBS buffer via ultrafiltration.

4 mg chemically synthesized peptide H3 (240-279) was ordered from GenScript Corporation.

**Modeling and MD Simulations of C-terminal Insertion of the H3 into DPPC Membrane**

Using TMHMM2.0^1^, we predicted the transmembrane region of H3 to span residues 283 to 306. We modeled this region using ESMFold, obtaining the helical structure of H3(283-306). Subsequently, by utilizing the previously obtained structures of H3(1-282) and H3(283-306), we performed multi-template modeling of the full-length H3 protein using MODELLER^2^. Next, we employed CHARMM-GUI’s^3^ Membrane Builder to construct the structure of H3 inserted into a 10 nm × 10 nm DPPC membrane, and simultaneously built a water box with a NaCl concentration of 0.15 M, generating input files (charmm36m force field and the TIP3P water model) suitable for MD simulations with GROMACS 2023.3^4^.

After energy minimization to reduce the maximum force below 1000 kJ/mol/nm, an NVT equilibration was conducted using the V-rescale thermostat method at 310K. This was followed by a 500 ns NpT production simulation at 310K using the V-rescale thermostat and the C-rescale barostat methods at 1 bar. During the 500 ns MD simulation, the time-dependent changes in the minimum distance between the helical region H3(240-273) and the DPPC membrane were calculated from the simulation trajectory using *gmx mindist*.

**Molecular Docking of H3 with HS**

A HS oligosaccharide with the structure -[IdoA2S-GlcNS6S-IdoA-GlcNS(3,6S)]_5_- was modeled using the CHARMM-GUI online website. The structure of H3(1-282) predicted by AF2 was refined through 500 ns of molecular dynamics simulation, and the most frequently appearing conformation was obtained through structural clustering. Specifically, by calculating the RMSD values of H3 in the MD trajectory, and then clustering using the built-in gmx cluster program, the gromos clustering method was adopted with a cut-off set to 0.2 nm. The top three conformations obtained were used for the subsequent docking process.

AutoDock Vina was called via the Chimera graphical interface to generate docking configuration files for HS and H3, with the center of each docking box set at the center of the GAGs binding motif, and the box size set to 8 nm. The energy_range was set to 3 kcal/mol, specifying that the energy difference from the current best conformation should be less than 3 kcal/mol. The docking results were ranked according to the energy scores generated by AutoDock Vina, and the best four conformations were selected.

**MD Simulations of the H3-HS Complex**

The docked H3-HS complex was uploaded to the CHARMM-GUI website for simulation system construction, utilizing the charmm36m force field and the TIP3P water model. A water box, 1 nm larger than the molecular boundaries of the complex, was constructed, and 0.15 M NaCl was used for charge balancing. The final output included structure and force field files suitable for GROMACS 2023.3 MD simulations. After energy minimization to reduce the maximum force below 1000 kJ/mol/nm, an NVT equilibration was conducted using the V-rescale thermostat method at 310K for 1 ns. This was followed by a 1000 ns NpT production simulation at 310K using the V-rescale thermostat and the C-rescale barostat methods at 1 bar. Three independent repetitions were performed for the production simulation.

The RMSD values of HS were calculated using the built-in gmx rms program in GROMACS, maintaining the structural overlap of H3. To analyze salt bridges between HS and HS in this simulation, a Python script was used to calculate the distance between the negatively charged oxygen atoms on the side chains of HS and the nitrogen atoms on the side chains of basic amino acids of H3 throughout the trajectory. A distance less than the set cutoff of 0.35 nm was considered indicative of salt bridge formation.

**Binding Free Energy of H3 with HS Using Umbrella Sampling in MD Simulations**

To calculate the stability of various binding conformations of H3-HS, the binding free energy was determined using the umbrella sampling method. The force field and solvent settings of the simulation system were consistent with the previously mentioned MD simulations, employing the CHARMM36m force field and TIP3 water model, and maintaining a simulation temperature of 310K. These simulations were conducted using GROMACS 2023.3. Specifically, for umbrella sampling, Steered Molecular Dynamics (SMD) simulations were first carried out. During this process, H3 was held fixed (using a restraining potential), and a harmonic potential was applied to the HS molecule. This potential facilitated the constant velocity stretching of HS (1 nm/ns) along the z-axis, moving it 5 nm until complete separation from H3 was achieved. The size of the simulation box was designed to ensure that neither molecule crossed the boundaries during stretching. The stretching process generated a series of reaction coordinates, defined by the distance between the centroids of HS and H3. Biasing potentials were applied to these coordinates for 10 ns MD simulations. The outputs of these simulations, in the form of potential energy distributions, were then analyzed using the Weighted Histogram Analysis Method (WHAM). This analysis produced a Potential of Mean Force (PMF) profile, describing the free energy landscape as a function of the reaction coordinate, thus allowing for the calculation of the binding free energy of H3-HS.

**Replica Exchange Molecular Dynamics Simulations**

To further investigate the conformational changes of HS and the helical domain, we conducted replica exchange MD simulations. The simulation system was set up using CHARMM-GUI, employing the same force field and water model as the aforementioned MD simulations, and 0.15 M NaCl was used for charge balancing. The temperature range was set from 310K to 387K, generating a total of 64 replicas with a swap probability of 20%. Each simulation had a duration of 1 µs, resulting in a cumulative simulation time of 64 µs for all replicas.

Simulations were executed using the MPI version of GROMACS 2023.3. Upon completion of the simulations, the replica at 310K was selected for further analysis. Using GROMACS built-in tools, 'distance' and 'rms', we calculated the nearest distance between HS and Mg(II), and the RMSD of HS throughout the simulation trajectory, respectively. Subsequently, the gmx sham program was used to generate a free energy landscape from these two sets of data. This landscape effectively illustrates the dynamic binding changes between HS and H3, as well as the interaction between HS and Mg(II).

***Poxviridae* Virus H3 Sequence Analysis**

A total of 66 available H3 sequences from *Poxviridae* viruses were obtained through a sequence search of the monkeypox virus H3 sequence using NCBI's BLAST^5^. After removing duplicate sequence data, multiple sequence alignment was performed using MEGA11^6^, with the ClustalW method^7^ employed for sequence alignment. The alignment results were uploaded to the WebLogo website^8^ for logo image generation. Acidic amino acids, basic amino acids, polar amino acids, and non-polar amino acids were represented in red, blue, green, and black, respectively. The font size in the logo corresponds to the probability of occurrence of the amino acid at that position, with larger fonts indicating greater conservation of the amino acid. The charge analysis of the amino acid sequences was conducted using the ProteinAnalysis function of the Biopython package. Surface charge maps of the H3 protein were generated using Chimera, and the charge distribution was calculated using APBS.

**AFM-SMFS Protein Unfolding Experiment**

The AFM cantilever/tip made of silicon nitride (MLCT-BIO-DC, Bruker Corp.) was used. The detailed protocol for AFM tip functionalization and protein immobilization on the glass coverslip can be found in references^9,10^. In short, the tip and glass coverslip were coated with the amino group by amino-silanization. N_3_ is functionalized on the surface from the reaction between ImSO_2_N_3_·HCl and –NH_2_. Then, a heterobifunctional DBCO-PEGn-Mal can be reacted and adds the Mal group. Next, the peptide GL-ELP_20_-C or C-ELP_20_-NGL was reacted to the maleimide via the cysteine, respectively. The long ELP_20_ serves as a spacer to avoid non-specific interaction between the tip and the surface as well as a signature for the single-molecule event. Finally, target protein H3 with C-terminal NGL sequence or GB1-Doc with N-terminal GL sequence can be site-specifically linked to the coverslip or tip by AEP, respectively.

Atomic force microscope (Nanowizard4, JPK) was used to acquire the force-extension curve. The D tip of the MLCT-Bio-DC cantilever was used. Its accurate spring constant was determined by a thermally-induced fluctuation method ^11^. Typically, the tip contacted the protein-immobilized surface for 100 *ms* under an indentation force of 350 pN to ensure a site-specifically interaction. Then, moving the tip up vertically at a constant velocity (1 µm/s), the polyprotein unfolded. Then, the tip moved to another place to repeat this cycle several thousands of times. As a result, a force-extension curve was obtained, which was analyzed using JPK data process analysis software.

**AFM-SMFS Unbinding Experiment of H3-HS on CHO-K1**

Force-extension based AFM measurement on model surfaces was performed in PBS buffer at room temperature using functionalized D tip of MLCT-Bio-DC cantilever (Bruker, nominal spring constant of 0.030 N/m and actual spring constants calculated using thermal tune). AFM (Nanowizard4, JPK) operated in the force mapping (contact) mode was used. CHO-K1 cells were cultured in a 37°C CO_2_ incubator for 24 hours prior to force spectroscopy experiments. The relative positioning of the AFM probe and the adherent cells was determined using an inverted microscope. Areas of 5 × 5 µm were scanned, ramp size set to 350 nm, and set point force of 300 pN with a contact time of 200 ms, with a resolution of 32 × 32 pixels.

**Flow Cytometry Experiment**

Protein binding to glycosaminoglycans (GAGs) expressed on the surface of Chinese Hamster Ovary K1 (CHO-K1) cells was assessed using flow cytometry, following the previously described methods. H3-eGFP, H3(uncharged)-eGFP, and eGFP proteins (5 μM) were incubated with 1 million cells (200 μL) at 4℃ for 20 minutes. Post-incubation, cell analysis was performed using the CytoFLEX flow cytometer (Beckman) to collect fluorescence signals in the FITC channel. A total of 15,000 events were collected using the CytExpert software (version 2.5; Beckman), with 10 sets of data collected in parallel. The data from the live-cell gating were analyzed using FlowJo software (version 10.6.2; Tree Star, San Carlos, CA).

The flow cytometry experiment to validate the inhibitory effects of inhibitors utilized the same equipment and methods as previously described. The final concentrations of H3-eGFP and the inhibitor were 2 μM and 10 μM, respectively, with a total volume of 600 μL. The control group contained only 2 μM H3-eGFP, with the volume made up with PBS. Data from five parallel experiments were collected, with each experiment gathering 15,000 events. Using FlowJo, the fluorescence intensity of the mean of FITC-area signal in the live cell gate was analyzed, normalizing the relative fluorescence intensity of the control group to 1.

To further verify the concentration-dependent inhibitory effect of AI-Poxblock 723, we conducted a concentration-dependent flow cytometry experiment. The final concentration of H3-eGFP remained at 2 μM, while 723 was tested at three final concentrations (100 μM, 50 μM, and 10 μM), with a total volume of 600 μL. The control group contained only 2 μM H3-eGFP without inhibitor. Data from five parallel experiments were collected, with 15,000 events recorded for each experiment. The mean fluorescence intensity in the FITC-area signal within the live cell gate was analyzed using FlowJo software, normalizing the relative fluorescence intensity of the control group to 1.

**Design of Inhibitors Targeting the H3 Helical Domain**

To design inhibitors targeting the helical domain of H3, hotspots information was first inputted into RFdiffusion. The selected residues for this purpose were 239, 242, 267, 266, 259 of H3, and the length of the inhibitor sequences was set to be 40-80 amino acids. This process generated 1000 backbone structures. After excluding single-stranded helical structures, the remaining 633 structures underwent sequence recovery using ProteinMPNN, producing 10 sequences per structure. All 6330 sequences were then subjected to structure prediction and scoring using AF2. Structures with an iPTM score greater than 0.9 and a pAE score less than 6 were selected, while sequences containing Cys were excluded to avoid the formation of disulfide bonds.

RMSD calculations were performed between the predicted structures and the RFdiffusion-designed structures, and sequences with an RMSD greater than 1.5 nm were excluded to ensure consistency between the predicted and designed structures. Additionally, solubility assessments were conducted for all sequences. Specifically, a solubility parameter was assigned to each of the 20 natural amino acids, and an average was calculated for the entire sequence. The sequences with the highest solubility were selected for expression and interaction testing.

**MD Simulations of H3 and its Inhibitors.**

The AF2-predicted structure of the H3-inhibitor complex was used for the MD simulations, utilizing the previously mentioned MD software and force field. The MD simulation was conducted at 310K for 500 ns. Subsequently, the RMSD of the simulation trajectory was calculated using the built-in 'rms' program of GROMACS to analyze the stability of the inhibitor-H3 binding.

**Circular Dichroism Experiments**

For circular dichroism (CD) experiments, AI-Poxblock723 and H3(240-279) were diluted to 0.15mg/mL and 0.1 mg/mL in 10 mM K-PO_4_ (pH 7.4) buffer, respectively. Spectra were acquired on a Chirascan V100 (Applied Photophysics). The data acquisition wavelength is set to 190-260 nm. All reported measurements were acquired within the linear range of the instrument.

**Bio-layer Interferometry (BLI) Binding Experiments**

BLI experiments were conducted using Octet HIS1K Biosensors on the Octet system. The buffer used throughout the experiments was uniform PBS. The tips were first pre-equilibrated in PBS solution for 10 minutes, followed by a 60 s baseline, 60 s loading, 100 s baseline, 100 s association, and 120 s dissociation steps. Baseline measurements of unloaded tips were subtracted from their matched measurement of the loaded tip. The inhibitor concentration was determined using the BCA method before loading and prepared at a concentration of 0.1 mg/mL for the loading solution. The mass concentration of H3(1-282) and H3(1-239) was also determined using the BCA method, and their molar concentrations were calculated. The purity of H3 is above 60%. Various concentration samples were then prepared using a serial dilution method. After baseline correction and curve smoothing, a global fit was performed to calculate the K_D_.

**Inhibition Test for Viral Infection**

Vero E6 cells were seeded at 1.5 × 10^4^ cells/well into 96-well plates and used the following day. AI-Poxblocks 3-fold serial diluted in maintenance medium were mixed with an equal volume of diluted live VACV or MPXV and then incubated at 37 °C for 1 hour. Medium from 96-well plates was aspirated, and the inhibitor-virus mixture was added (100 μl/well), then the Vero E6 cells were incubated at 37 °C for about 16 hours. Then cells were fixed with 4% paraformaldehyde solution, permeabilized with Perm/Wash buffer (BD Biosciences) containing 0.1% Triton X-100, incubated with the HRP-conjugated anti-VACV polyclonal antibodies (Invitrogen) diluted in the Perm/Wash buffer at room temperature for 2 hours.

The reactions were developed with KPL TrueBlue Peroxidase substrates (Seracare Life Sciences). The numbers of infected foci were calculated using an EliSpot reader (Cellular Technology Ltd). The 50% inhibitory concentration (IC50) was calculated using GraphPad Prism software using the log (inhibitor) vs. normalized response-variable slope (four parameters) model. Experiments with live MPXV were performed in a Biosafety Level 3 (BSL-3) facility following standard biosafety practices.

**Animal Experiments**

Female BALB/c mice, 6-8 weeks old, were obtained from GemPharmatech Co., Ltd (Guangdong, China). After arrival, all mice were acclimated for 3 days and then randomly assigned to experimental groups. For challenge, mice were anesthetized by intraperitoneal injection of 1.25% tribromoethanol solution with a dosage of 20 μL/g. Intranasal (i.n.) infections were performed by introduction of 50 μL of virus into one nostril. Mock-infected control animals were similarly inoculated with an equivalent volume of diluent. After infection, mice were given intraperitoneally with single dose of 10 mg/kg of AI-Poxblock723 or PBS. animals were euthanized 4 days after infection for the determination of pulmonary viral titers.

**Animal Ethics Statement**

Mice were housed in independent ventilation cages and utilized at five mice per treatment group. All animals were given food and water ad libitum throughout all experiments. All efforts were made to minimize animal suffering and to reduce the number of animals used. All animal procedures were approved by Shenzhen Third People’s Hospital’s Institutional Animal Care and Use Committee prior to the initiation of studies.

**Infectious Virus Titration and Viral Load Determination.**

The lungs from BALB/c mice were homogenized in 1 mL phosphate buffer solution (PBS) with 3 mm zirconium beads using a tissue homogenizer (Omni, Bead Ruptor 24 Elite). After two cycles of freeze-thaw, the lung tissue homogenates were centrifuged to remove tissue debris. Supernatants were transferred into fresh tubes. Infectious viral titers were determined using focus forming assay. Briefly, the Vero E6 cells were seeded in 96 well plates and incubated overnight. Ten-fold serial dilutions of the MPXV were prepared in the maintenance medium. The Vero E6 cells were inoculated with 100-fold diluted supernatants (100 μL/well). After 1 h of adsorption, the samples were removed and cells were washed once with PBS. Then 100 μL/well maintenance medium was added. After 18 h incubation, the medium was aspirated and the cells were fixed with 4% paraformaldehyde solution for 30 min, permeabilized with Perm/Wash buffer (BD Biosciences) containing 0.1% Triton X-100, incubated with the HRP-conjugated anti-VACV polyclonal antibodies (Invitrogen) at room temperature for 2 hours. The reactions were developed with KPL TrueBlue Peroxidase substrates (Seracare Life Sciences). The number of virus foci was counted using an ELISpot reader (Cellular Technology Ltd.).

MPXV viral loads (genome DNA copies) were determined using quantitative polymerase chain reaction (qPCR). Briefly, MPXV DNA in tissue homogenate supernatants was extracted using a nucleic acid extraction instrument (DaAn Gene, Smart 32) combined with Daan's extraction kit according to the manufacturer’s instructions. The MPXV DNA copies were determined using SYBR Green Premix Pro Taq HS qPCR Kit (Accurate Biotech) with the following primers: forward primer 5’-TTT ATT CAA CAT GTA CTG TAC CCA C-3’, reverse primer 5’- TTT CTT GCA TGG ATT TTC GTA TTT C -3’. MPXV B6R plasmid (Sangon Biotech) was serially diluted and performed to generate the standard curve.

**Supplementary Text**

**Protein Sequences:**

**AFM Experiment**

His6-Coh-GB1-GB1-H3(1-282)-strep tag Ⅱ-NGL

MRGSHHHHHHGSMGTALTDRGMTYDLDPKDGSSAATKPVLEVTKKVFDTAADAAGQTVTVEFKVSGAEGKYATTGYHIYWDERLEVVATKTGAYAKKGAALEDSSLAKAENNGNGVFVASGADDDFGADGVMWTVELKVPADAKAGDVYPIDVAYQWDPSKGDLFTDNKDSAQGKLMQAYFFTQGIKSSSNPSTDEYLVKANATYADGYIAIKAGEPRSMDTYKLILNGKTLKGETTTEAVDAATAEKVFKQYANDNGVDGEWTYDDATKTFTGTERSMDTYKLILNGKTLKGETTTEAVDAATAEKVFKQYANDNGVDGEWTYDDATKTFTGTERSMAAVKTPVIVVPVIDRPPSETFPNVHEHINDQKFDDVKDNEVMQEKRDVVIVNDDPDHYKDYVFIQWTGGNIRDDDKYTHFFSGFCNTMCTEETKRNIARHLALWDSKFFTELENKNVEYVVIIENDNVIEDITFLRPVLKAIHDKKIDILQMREIITGNKVKTELVIDKDHAIFTYTGGYDVSLSAYIIRVTTALNIVDEIIKSGGLSSGFYFEIARIENEMKINRQIMDNSAKYVEHDPRLVAEHRFETMKPNFWSRIGTVAAKRYPGVMYTFTTPLISFRSWSHPQFEKGSNGL

**H3 unbinding on CHO-K1 cells**

The following proteins were obtained by cleaving his6-TF-tev-POI using TEV protease.

H3(1-282)-strep tag Ⅱ-NGL

GGSMAAVKTPVIVVPVIDRPPSETFPNVHEHINDQKFDDVKDNEVMQEKRDVVIVNDDPDHYKDYVFIQWTGGNIRDDDKYTHFFSGFCNTMCTEETKRNIARHLALWDSKFFTELENKNVEYVVIIENDNVIEDITFLRPVLKAIHDKKIDILQMREIITGNKVKTELVIDKDHAIFTYTGGYDVSLSAYIIRVTTALNIVDEIIKSGGLSSGFYFEIARIENEMKINRQIMDNSAKYVEHDPRLVAEHRFETMKPNFWSRIGTVAAKRYPGVMYTFTTPLISFRSWSHPQFEKGSNGL

H3(uncharged)-strep tag Ⅱ-NGL

GGSMAAVKTPVIVVPVIDRPPSETFPNVHEHINDQKFDDVKDNEVMQEKRDVVIVNDDPDHYKDYVFIQWTGGNIRDDDKYTHFFSGFCNTMCTEETKRNIARHLALWDSKFFTELENKNVEYVVIIENDNVIEDITFLRPVLKAIHDKKIDILQMREIITGNKVKTELVIDKDHAIFTYTGGYDVSLSAYIIRVTTALNIVDEIIKSGGLSSGFYFEIARIENEMKINRQIMDNSAKYVEHDPSLVAESSFETMSPNFWSSIGTVAASSYPGVMYTFTTPLISFRSWSHPQFEKGSNGL

**Flow Cytometry Experiment**

Proteins were obtained by cleaving His6-TF-tev-POI using TEV protease.

H3(1-282)-eGFP- strep tag Ⅱ

GGSMAAVKTPVIVVPVIDRPPSETFPNVHEHINDQKFDDVKDNEVMQEKRDVVIVNDDPDHYKDYVFIQWTGGNIRDDDKYTHFFSGFCNTMCTEETKRNIARHLALWDSKFFTELENKNVEYVVIIENDNVIEDITFLRPVLKAIHDKKIDILQMREIITGNKVKTELVIDKDHAIFTYTGGYDVSLSAYIIRVTTALNIVDEIIKSGGLSSGFYFEIARIENEMKINRQIMDNSAKYVEHDPRLVAEHRFETMKPNFWSRIGTVAAKRYPGVMYTFTTPLISFRSPAPAPAPMVSKGEELFTGVVPILVELDGDVNGHKFSVSGEGEGDATYGKLTLKFICTTGKLPVPWPTLVTTLTYGVQCFSRYPDHMKQHDFFKSAMPEGYVQERTIFFKDDGNYKTRAEVKFEGDTLVNRIELKGIDFKEDGNILGHKLEYNYNSHNVYIMADKQKNGIKVNFKIRHNIEDGSVQLADHYQQNTPIGDGPVLLPDNHYLSTQSALSKDPNEKRDHMVLLEFVTAAGITLGMDELYK-RSWSHPQFEK

H3(uncharged)-eGFP- strep tag Ⅱ

GGSMAAVKTPVIVVPVIDRPPSETFPNVHEHINDQKFDDVKDNEVMQEKRDVVIVNDDPDHYKDYVFIQWTGGNIRDDDKYTHFFSGFCNTMCTEETKRNIARHLALWDSKFFTELENKNVEYVVIIENDNVIEDITFLRPVLKAIHDKKIDILQMREIITGNKVKTELVIDKDHAIFTYTGGYDVSLSAYIIRVTTALNIVDEIIKSGGLSSGFYFEIARIENEMKINRQIMDNSAKYVEHDPSLVAESSFETMSPNFWSSIGTVAASSYPGVMYTFTTPLISFRSPAPAPAPMVSKGEELFTGVVPILVELDGDVNGHKFSVSGEGEGDATYGKLTLKFICTTGKLPVPWPTLVTTLTYGVQCFSRYPDHMKQHDFFKSAMPEGYVQERTIFFKDDGNYKTRAEVKFEGDTLVNRIELKGIDFKEDGNILGHKLEYNYNSHNVYIMADKQKNGIKVNFKIRHNIEDGSVQLADHYQQNTPIGDGPVLLPDNHYLSTQSALSKDPNEKRDHMVLLEFVTAAGITLGMDELYK-RSWSHPQFEK

eGFP- strep tag Ⅱ

GGSMVSKGEELFTGVVPILVELDGDVNGHKFSVSGEGEGDATYGKLTLKFICTTGKLPVPWPTLVTTLTYGVQCFSRYPDHMKQHDFFKSAMPEGYVQERTIFFKDDGNYKTRAEVKFEGDTLVNRIELKGIDFKEDGNILGHKLEYNYNSHNVYIMADKQKNGIKVNFKIRHNIEDGSVQLADHYQQNTPIGDGPVLLPDNHYLSTQSALSKDPNEKRDHMVLLEFVTAAGITLGMDELYK-RSWSHPQFEK

**Inhibitor Design**

AI-Poxblock602-strep tag Ⅱ-His8:

MSAEAAKLAKKAVEDPSYAEELLKKDAESSTEERAAIANALLAERRRNPEKAQKMIEKAAKIAFEDAKKELEEEEKKKAERSGGGGSGGGGSWSHPQFEKGGSHHHHHHHH

MW: 12.14 kDa

AI-Poxblock614-strep tag Ⅱ-His8:

MGLSKLNELKAKANALGKEARAALDAGDFEKAKEKILAGAKATKEAGELSGNKAMIEDGEKAEEVAEEIVKIAKEERAKERSGGGGSGGGGSWSHPQFEKGGSHHHHHHHH

MW: 11.68 kDa

AI-Poxblock723-strep tag Ⅱ-His8:

MGLKKLQELKKKANELGKKAKEALDKGDFEKAKEYIKKGAEATKEYGELSGNKAAIEDGEKAEEVAEEIVKIAKEEKAKERSGGGGSGGGGSWSHPQFEKGGSHHHHHHHH

MW: 12.05 kDa

AI-Poxblock761-strep tag Ⅱ-His8:

MSEFLESLKKAEELRKEVRELMEKGKEEADKLYKEGKEEEAAKVFLETAKEAEPIAEEALKLLMERSGGGGSGGGGSWSHPQFEKGGSHHHHHHHH

MW: 10.74 kDa

The following proteins were obtained by cleaving TF-tev-POI-strep with TEV protease. To avoid the influence of the His tag from the pCold vector on BLI assays, site-directed mutagenesis via PCR was employed to remove the His tag present in the vector.

H3(1-282)-strep tag Ⅱ-NGL

GGSMAAVKTPVIVVPVIDRPPSETFPNVHEHINDQKFDDVKDNEVMQEKRDVVIVNDDPDHYKDYVFIQWTGGNIRDDDKYTHFFSGFCNTMCTEETKRNIARHLALWDSKFFTELENKNVEYVVIIENDNVIEDITFLRPVLKAIHDKKIDILQMREIITGNKVKTELVIDKDHAIFTYTGGYDVSLSAYIIRVTTALNIVDEIIKSGGLSSGFYFEIARIENEMKINRQIMDNSAKYVEHDPRLVAEHRFETMKPNFWSRIGTVAAKRYPGVMYTFTTPLISFRSWSHPQFEKGSNGL

H3(uncharged)-strep tag Ⅱ-NGL

GGSMAAVKTPVIVVPVIDRPPSETFPNVHEHINDQKFDDVKDNEVMQEKRDVVIVNDDPDHYKDYVFIQWTGGNIRDDDKYTHFFSGFCNTMCTEETKRNIARHLALWDSKFFTELENKNVEYVVIIENDNVIEDITFLRPVLKAIHDKKIDILQMREIITGNKVKTELVIDKDHAIFTYTGGYDVSLSAYIIRVTTALNIVDEIIKSGGLSSGFYFEIARIENEMKINRQIMDNSAKYVEHDPSLVAESSFETMSPNFWSSIGTVAASSYPGVMYTFTTPLISFRSWSHPQFEKGSNGL

**Supplementary Figures**

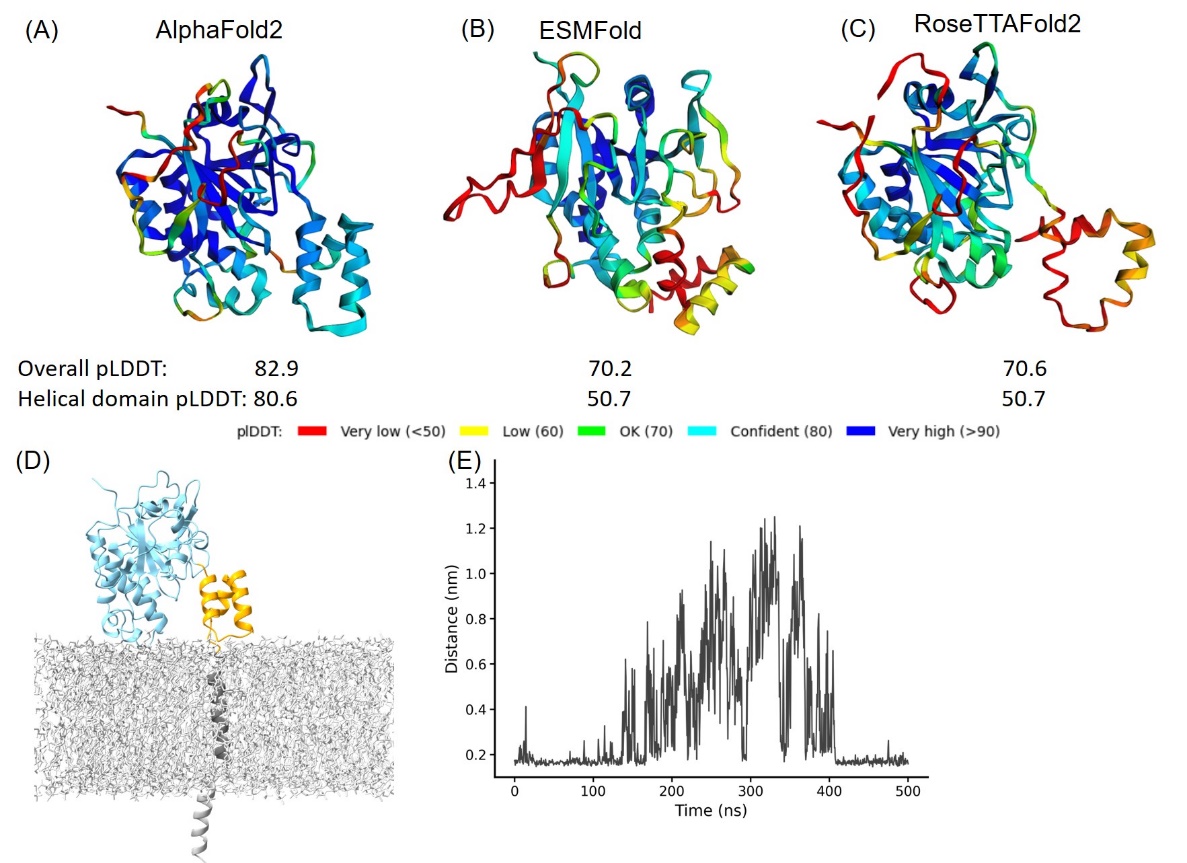

**Fig. S1 Structural Prediction Protein-Membrane Modelling of Monkeypox Virus H3 Protein.** (A)-(C) In the protein cartoon diagram, the colors represent the pLDDT scores output by AF2, ESMFold and RoseTTAFold2 respectively, with blue indicating a higher prediction confidence. For the AF2 result, the helical domain is displayed in blue, indicating a region where the pLDDT score is greater than 80, signifying a high confidence in structural prediction. (D) Modelling of H3 with DPPC membrane. The blue represents H3(1-239), the yellow represents H3(240-282), the black corresponds to the transmembrane region H3(283-306), and the gray molecules represent DPPC. (E) The minimal distance between helical domain and DPPC membrane during the 500 ns MD simulations.

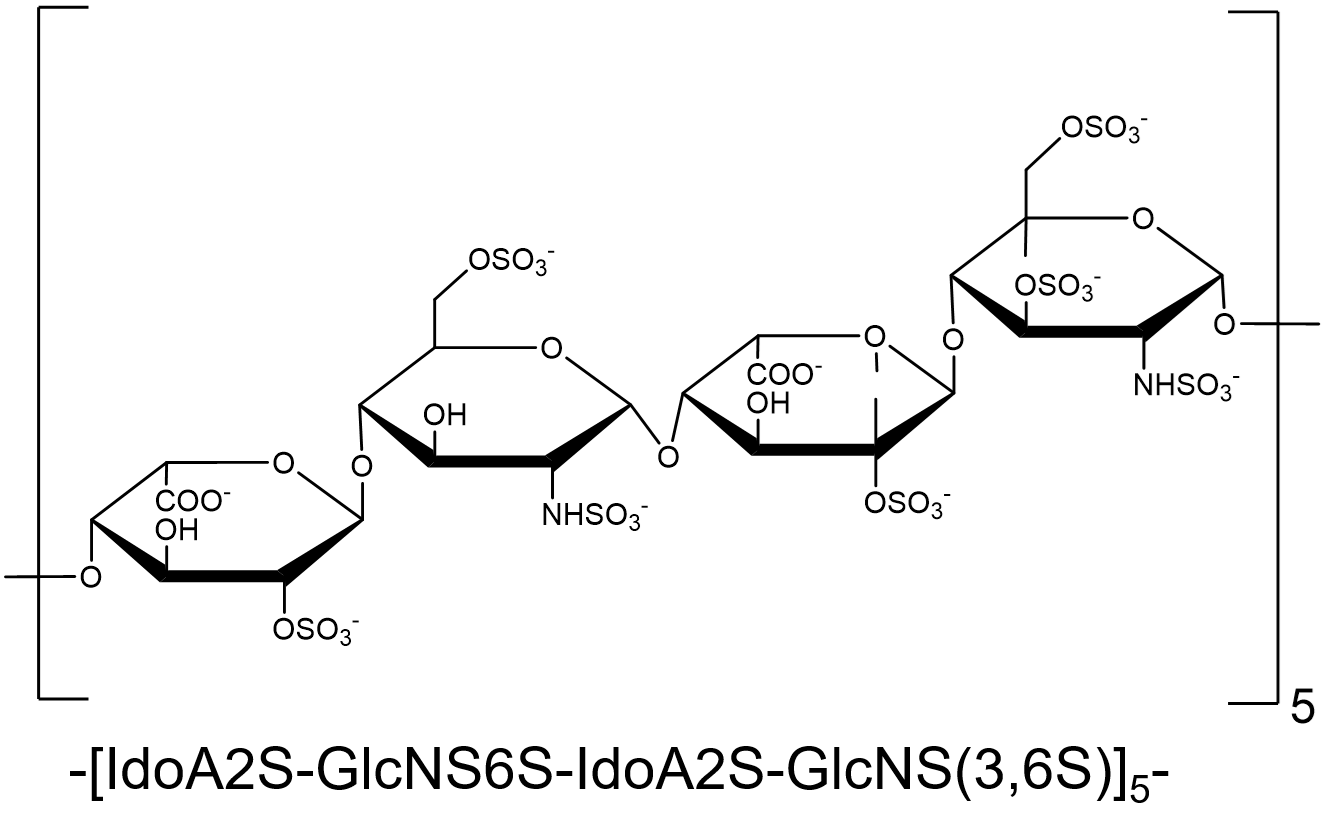

**Fig. S2 Structural Formula of HS.** The composition of HS used in the main text consists of -[IdoA2S-GlcNS6S-IdoA-GlcNS(3,6S)]_5_-, containing a total of 20 repeating monosaccharide units.

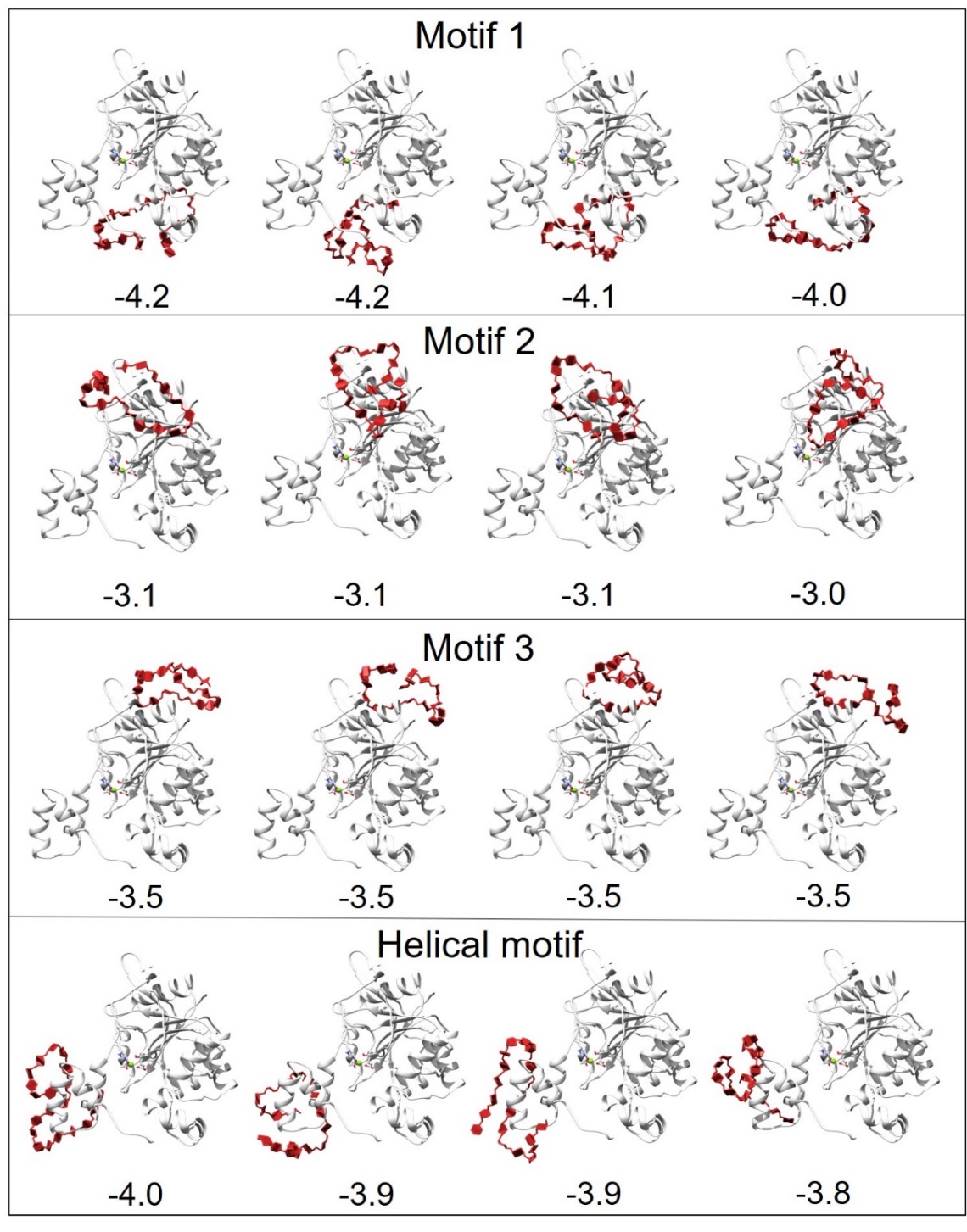

**Fig. S3 Docking Results of H3 with HS.** The docking area of HS was confined near different motifs by setting the docking box. The figure displays the docking results for four motifs, with the top four docking outcomes in each motif area selected based on the best scores (kCal/mol) from AutoDock Vina for subsequent MD simulations.

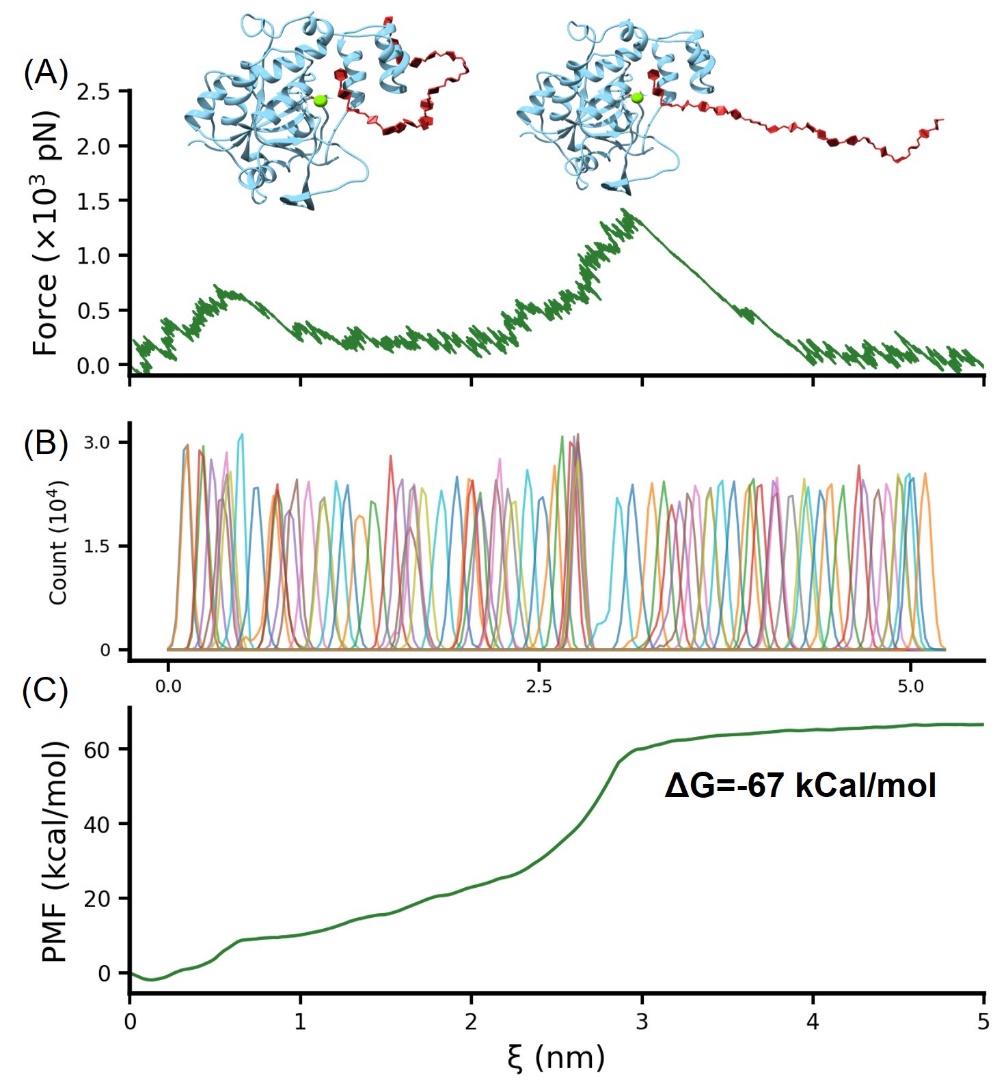

**Fig. S4 Umbrella Sampling Calculation of the Binding Free Energy of HS-H3.** The conformation of HS binding simultaneously to the helical domain and Mg^2+^, obtained from replica exchange simulations, was further subjected to umbrella sampling to calculate the binding free energy. The stretching process is illustrated in (A), with the distribution of the reaction coordinate shown in (B). (C) presents the PMF (Potential of Mean Force) profile obtained after calculations using the Weighted Histogram Analysis Method (WHAM).

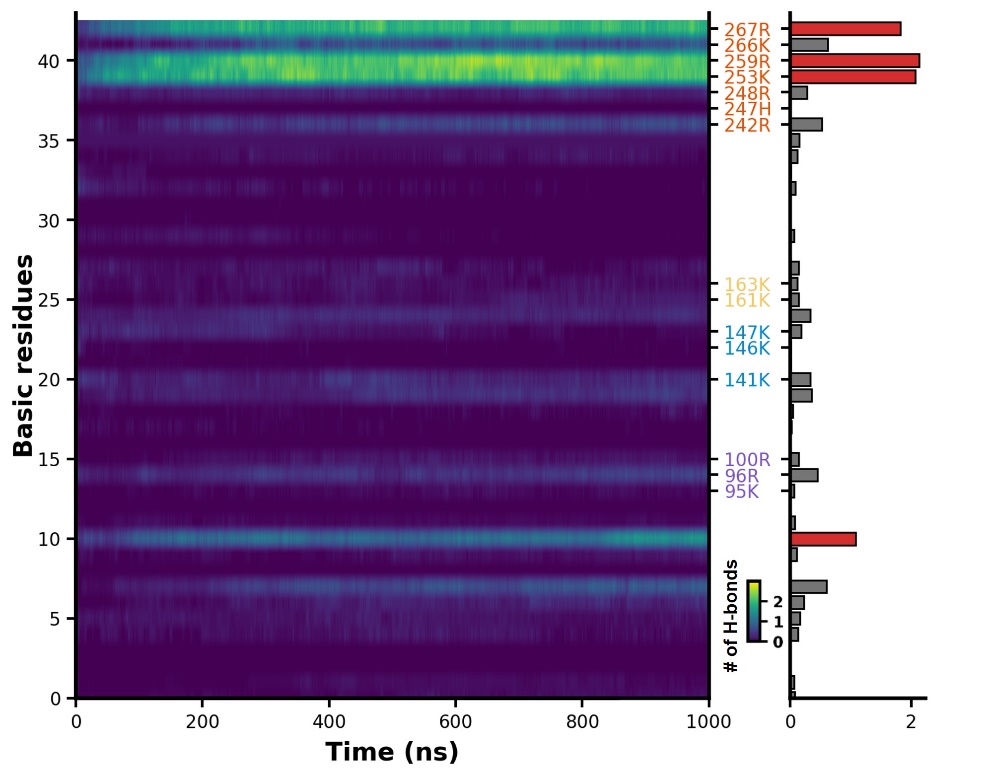

**Fig. S5 Evolution of Salt Bridge Formation in HS During MD Simulations.** This represents the variation in the number of salt bridges formed between HS and all basic amino acid side chain of H3 as the simulation progresses. The lighter areas indicate the formation of a greater number of salt bridges.

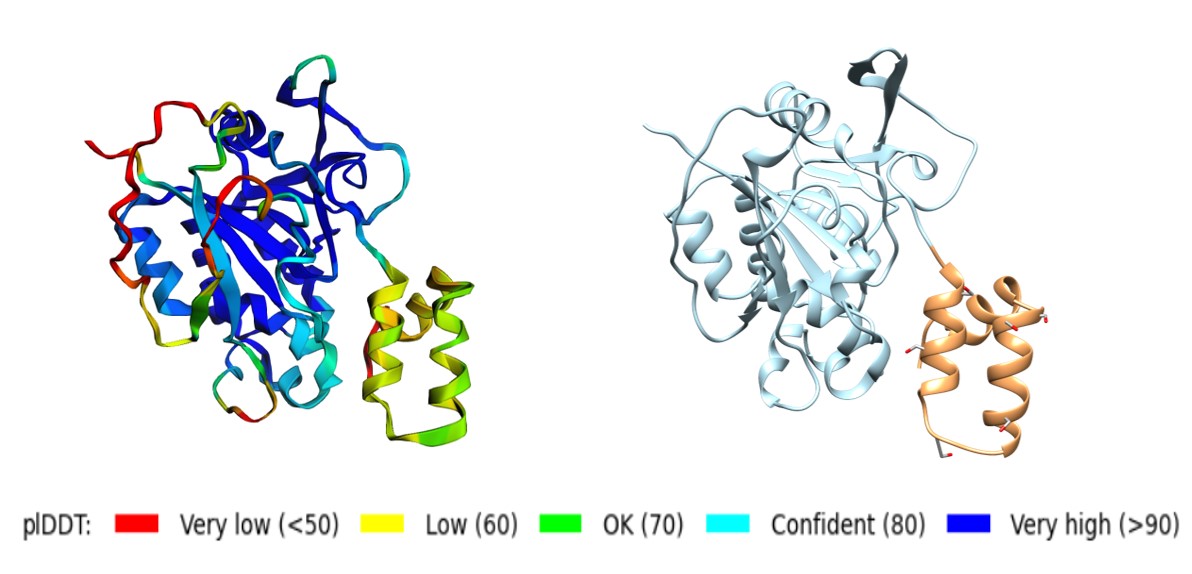

**Fig. S6** **Structure Prediction of H3 (uncharged) by AF2.** The left panel shows the pLDDT scores derived from the AF2 prediction. In the right panel, the blue region corresponds to H3(1-239), and the yellow region represents H3(240-282). All basic amino acids in H3(240-282) have been mutated to serine.

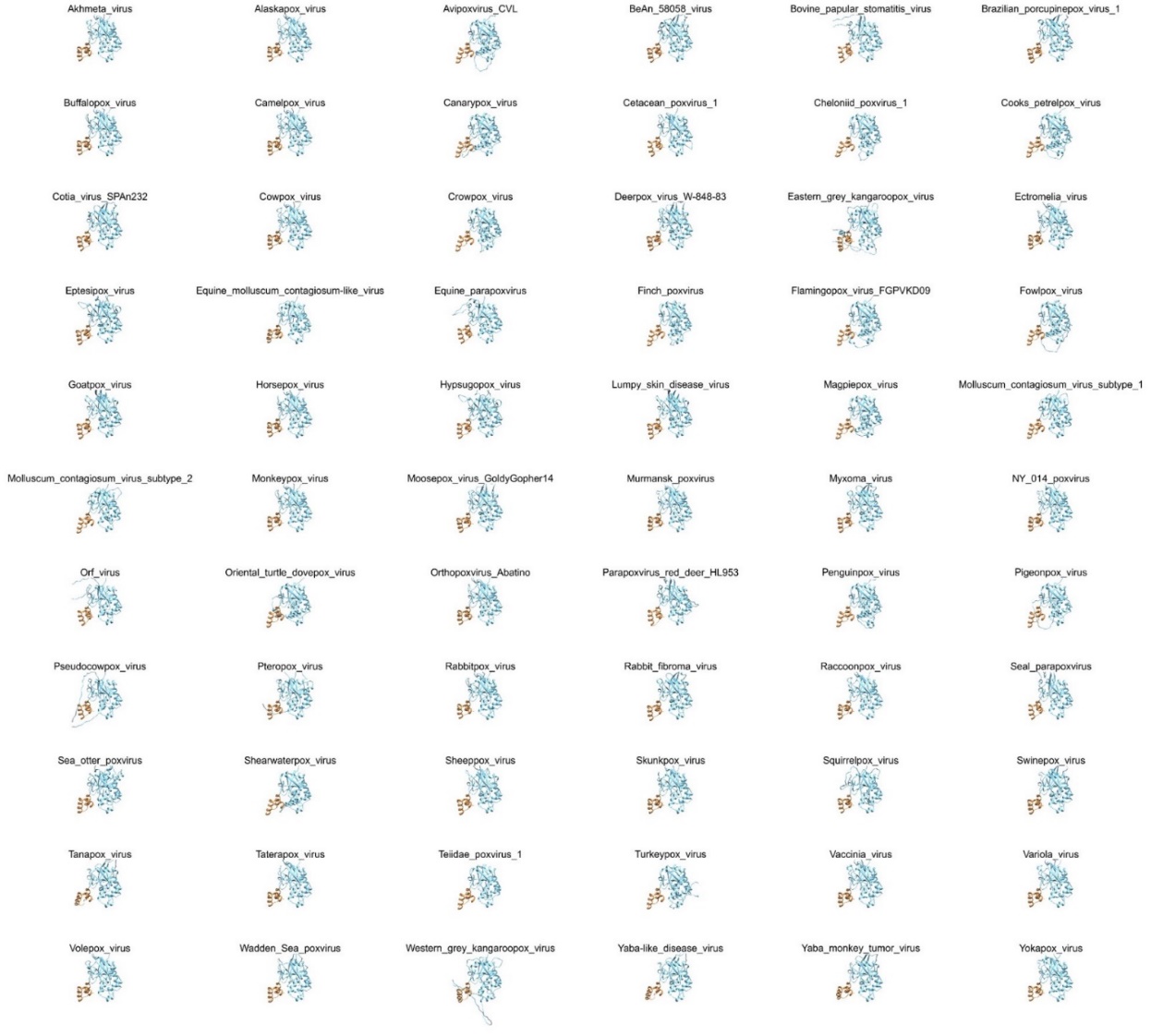
**Fig. S7 AF2 Structural Prediction of 66 *Poxviridae* Virus H3 Proteins.** The yellow portion represents the helical domain, while the blue portion signifies the main body of H3.

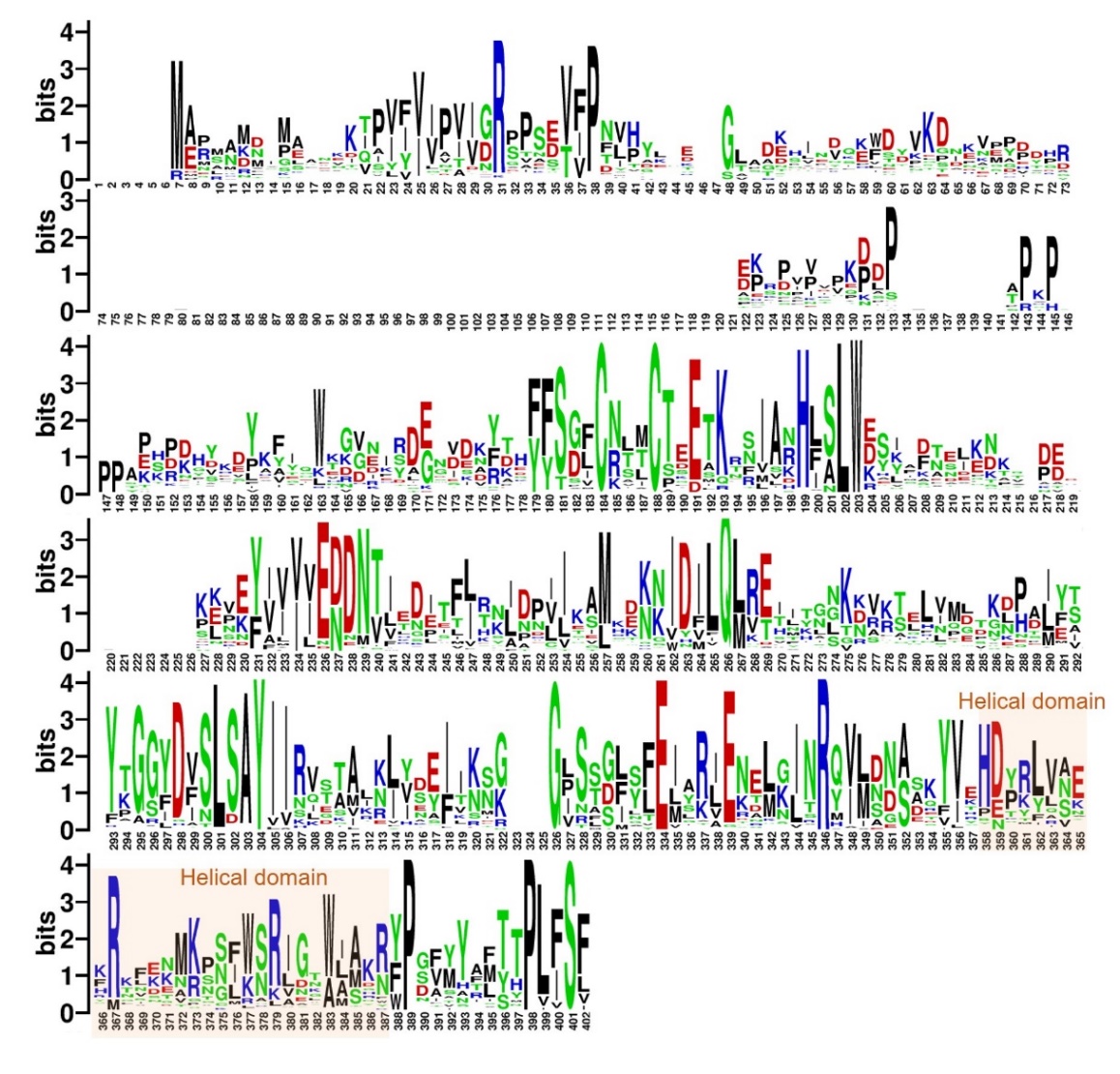

**Fig. S8 Analysis Results of Full Sequence Analysis for 66 *Poxviridae* Virus H3 Proteins.** The logo plot illustrates the probability of amino acid occurrence within the H3 sequences, where larger font sizes indicate a higher occurrence probability, denoting greater conservation. Acidic amino acids, basic amino acids, polar amino acids, and non-polar amino acids are represented in red, blue, green, and black, respectively.

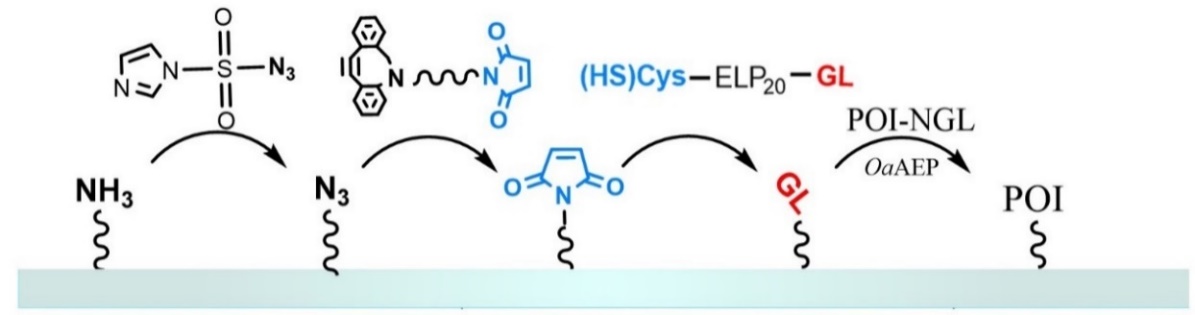

**Fig. S9 Protein Immobilization in AFM-SMFS Experiments.** Peptide GL-ELP_20_-Cys is modified on a glass substrate (or AFM tip) that has been functionalized with maleimide through a Michael addition reaction. The N-terminus GL can undergo an enzymatic ligation reaction with the C-terminus NGL of the target protein in the presence of ligase *Oa*AEP1, thereby immobilizing the target protein, such as Coh-GB1-GB1-H3-NGL and GL-GB1-Doc, on the substrate or AFM tip.

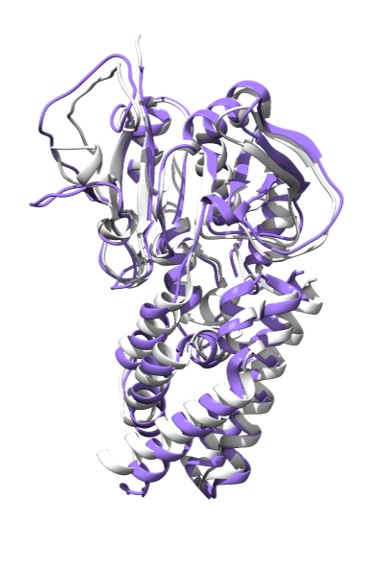

**Fig. S10 Artificial Design of H3 Inhibitors.** The gray structure represents the protein backbone structure designed by RFdiffusion, while the purple structure is the sequence generated by ProteinMPNN based on that backbone, followed by complex prediction results from AF2. The two structures align well, with an RMSD less than 1.5 nm.

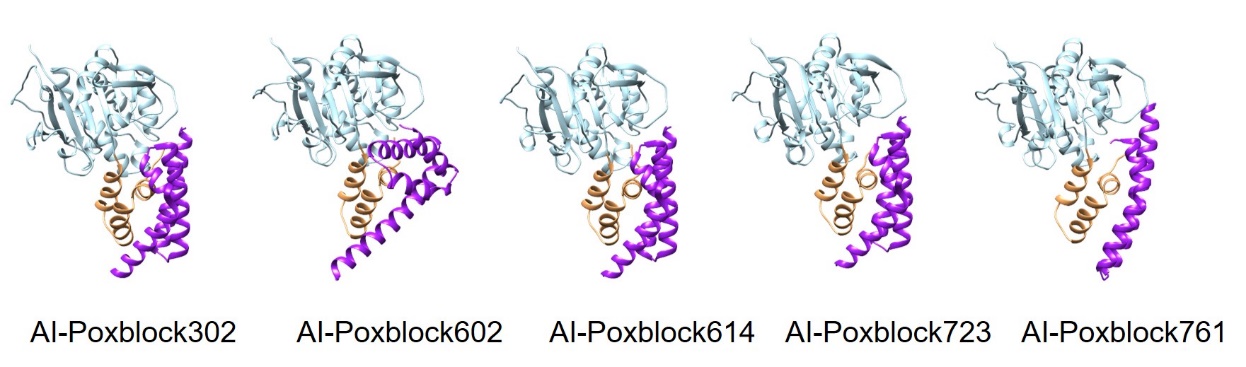

**Fig. S11 AF2-predicted Structures of the H3 and Inhibitor Complexes.** From left to right are AI-Poxblock302, AI-Poxblock602, AI-Poxblock614, AI-Poxblock723, and AI-Poxblock761. All inhibitors are depicted in purple, and the helical domain of H3 are marked in yellow.

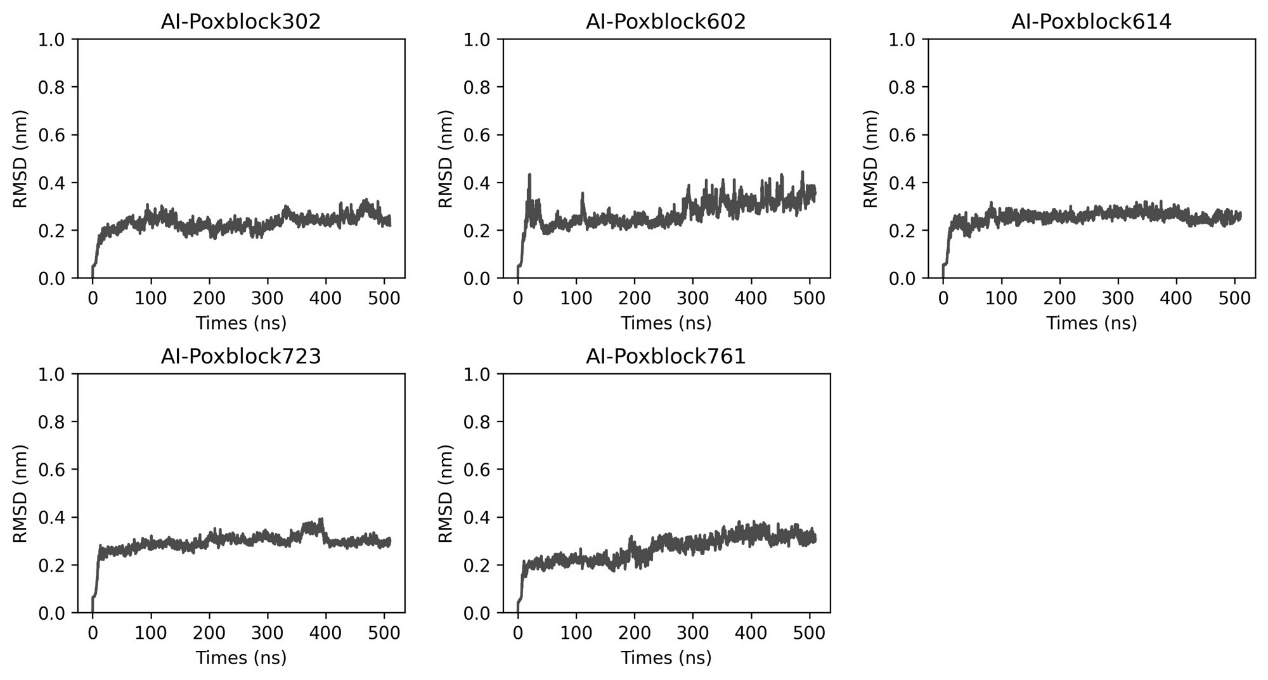

**Fig. S12 MD Simulations of H3-Inhibitor Complex.** The H3 and Inhibitor complex underwent a 500 ns MD simulation, resulting in the variation of RMSD (nm) over time (ns).

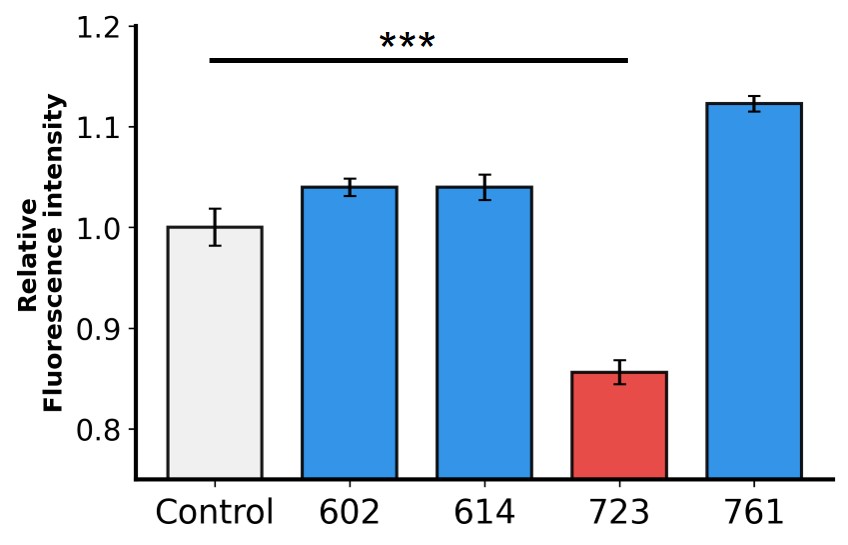

**Fig. S13 Flow Cytometry Analysis Demonstrates the Efficacy of the Inhibitors.** The graph shows relative fluorescence intensity, indicating that in the presence of 10 μM AI-Poxblock723, the binding of H3(1-282)-eGFP to CHO-K1 cells is reduced compared to the control group (gray).

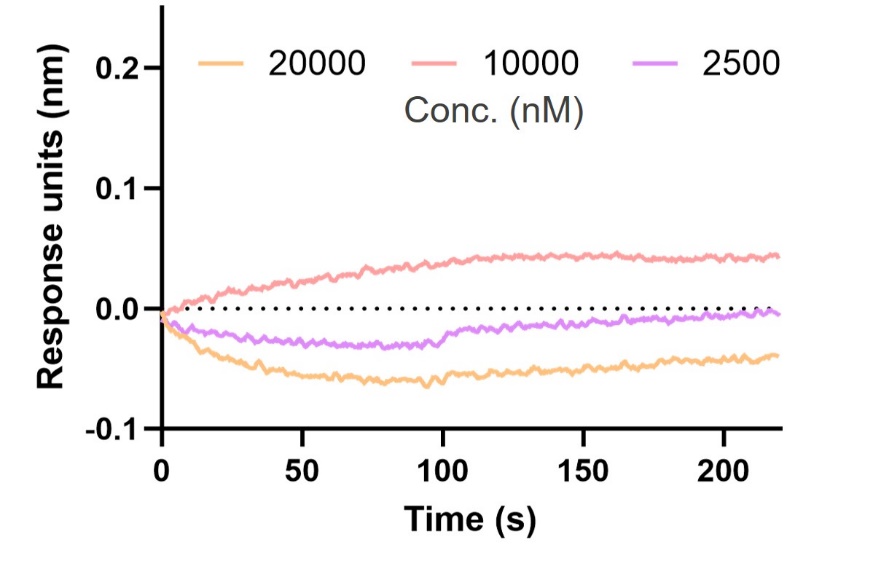

**Fig. S14 AI-Poxblock723 and H3(1-239) BLI Experiment.** BLI curves of three concentrations of H3(1-239) (20000 nM, 10000 nM, and 2500 nM) with AI-Poxblock723 loaded on the sensor. The curves could not be fitted to obtain the k_D_ value.

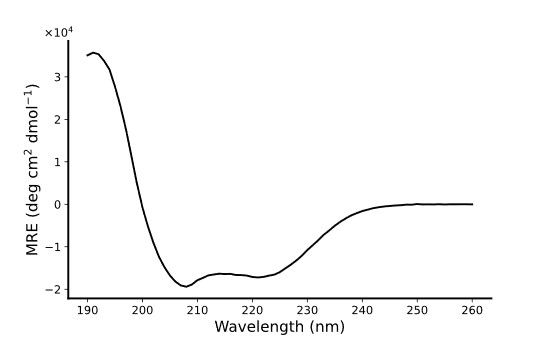

**Fig. S15 CD Spectroscopy of AI-Poxblock723.** CD spectroscopy verifies the designed inhibitor with a α-helical structure.

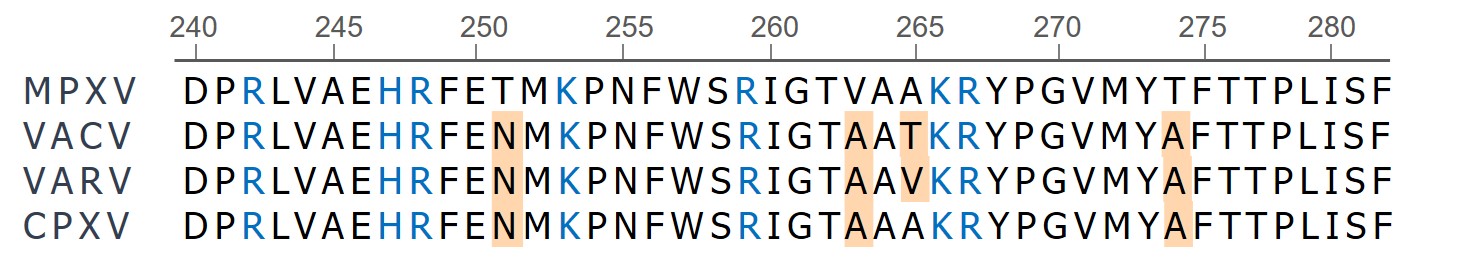

**Fig. S16 Sequence Comparison of the H3 Helical Domain among Representative *orthopoxvirus* strains MPXV, VACV, VARV, and CPXV.** Basic amino acid residues are highlighted in blue, and differences relative to the MPXV sequence are shaded in yellow.

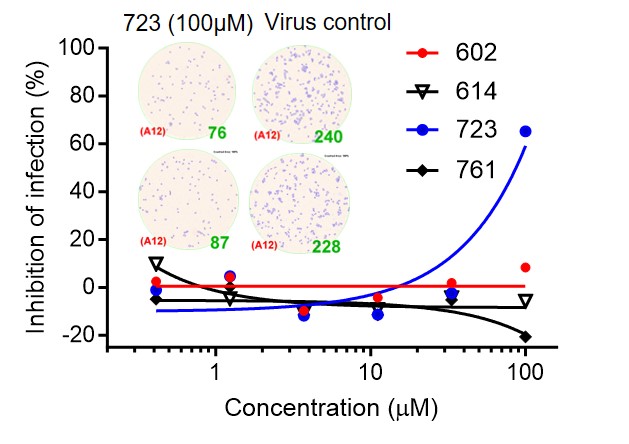

**Fig. S17 The Inhibitory Effect of Inhibitors on VACV Infection of Vero E6 Cells.** Virus-infected foci and counts were shown as inset.

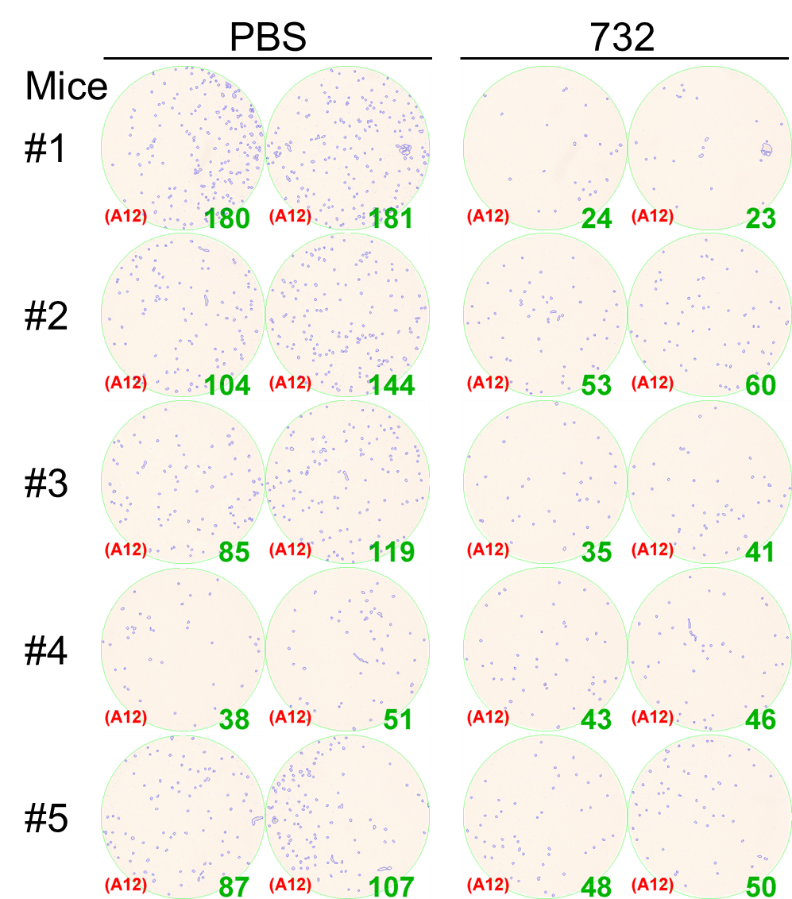

**Fig. S18 Titration of the MPXV Infectious Particles in Mice Lungs 4 Days Post-Infection.** Virus-infected foci and counts were shown as inset.

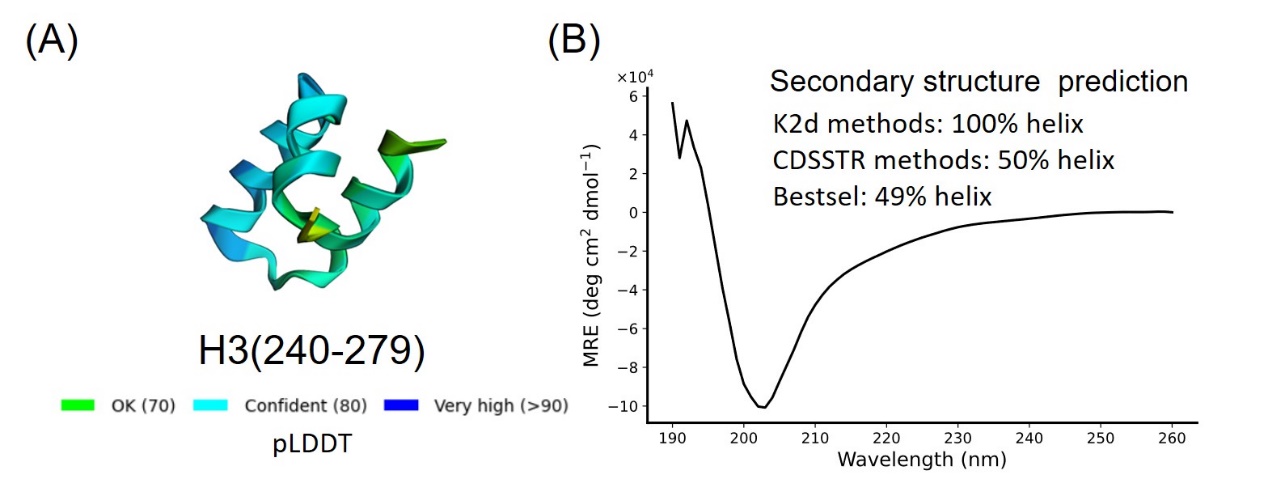

**Fig. S19 Secondary Structure Characterization of H3(240-279).** (A) AlphaFold2 prediction result of H3(240-279). (B) CD results and secondary structure prediction^12,13^ of H3(240-279).

**Supplementary Videos:**

Supplementary Video 1: MD simulation trajectories of H3 with DPPC membrane.

Supplementary Video 2: MD simulation trajectories of HS binding to H3 Domain 1 and the Helical domain.

Supplementary Video 3: SMD simulation trajectories of HS dissociating from H3 in umbrella sampling.

Supplementary Video 4: Trajectories of H3 and HS interactions in replica exchange MD simulations (REMD).

Supplementary Video 5: MD simulation trajectories of the H3-H3 complex after removal of the Mg (II) from H3.

Supplementary Video 6: MD simulation trajectories of the H3-H3 complex after mutating the positively charged amino acids on the helical domain of H3 to serine.

Supplementary Video 7: MD simulation trajectories of the AI-Poxblock723 complex.
